# Dual-patterned pluripotent stem cells self-organize into a human embryo model with extended anterior-posterior patterning

**DOI:** 10.1101/2025.09.25.678678

**Authors:** Zukai Liu, Chengxiang Qiu, Connor A. Kubo, Stella Xu, Riza M. Daza, Eva Nichols, Wei Yang, Anh Vo, Mary B. O’Neill, Choli Lee, Jay Shendure, Nobuhiko Hamazaki

**Affiliations:** Department of Genome Sciences, University of Washington, Seattle, WA, USA; Institute of Stem Cell and Regenerative Medicine, University of Washington, Seattle, WA, USA; Seattle Hub for Synthetic Biology, Seattle, WA, USA; Department of Molecular and Systems Biology, Dartmouth College, Hanover, NH, USA; Brotman Baty Institute for Precision Medicine, University of Washington, Seattle, WA, USA; Allen Discovery Center for Cell Lineage Tracing, Seattle, WA, USA; Howard Hughes Medical Institute, Seattle, WA, USA; Departments of Obstetrics & Gynecology, University of Washington, Seattle, WA, USA

## Abstract

Human gastruloids are a powerful class of stem cell-derived models that recapitulate key features of early embryonic development, including symmetry breaking and the emergence of three germ layers^1–3^. However, they lack anterior embryonic structures and coordinated axial organization^4–6^. To address this limitation, we pre-patterned human pluripotent stem cells (hPSCs) by exposing them to either anterior (FGF2) or posterior (CHIR99021 [CHIR] & retinoic acid [RA]) cues. Upon mixing, these dual-patterned hPSCs interacted and self-organized into elongated structures with both anterior and posterior features—which we term anterior-posterior (AP) human gastruloids. Anteriorly pre-treated cells robustly intercalated into posteriorly pre-treated cells, collectively giving rise to a continuum of neural tissues—including a brain-like domain, a neural tube-like structure, and neuro-mesodermal progenitors (NMPs)—with segmented somites arrayed bilaterally. Single cell RNA sequencing (scRNA-seq) revealed that human AP gastruloids contain cell types resembling the midbrain-hindbrain boundary (MHB), regionalized hindbrain structures (*i.e.* rhombomeres 1–8), regionalized neural crest (*i.e.* cranial, vagal, trunk)^7,8^ and head mesoderm. Transcriptomic comparisons to primate embryos revealed that human AP gastruloids most closely resemble Carnegie stage 11 (CS11) embryos. While they lack a notochord and full dorsal-ventral polarity, human AP gastruloids recapitulate key spatial and temporal features of early neurulation and somitogenesis. Perturbation of folic acid metabolism or rho-associated kinase (ROCK) signaling induced spinal cord defects, phenocopying aspects of spina bifida and other neural tube defects, highlighting this model’s potential for studying congenital disorders^9^. AP gastruloids may serve as a simple, robust, scalable platform for modeling coordinated human AP body axis development. More broadly, our results suggest that controlled interactions between differentially prepatterned progenitors can initiate self-organization of complex body axis features. The “pattern-and-mix” strategy may serve as a generalizable framework for assembling spatially organized stem cell models of mammalian development.

## INTRODUCTION

Human embryonic development is a highly orchestrated process that requires the precise coordination of molecular and cellular events across space and time. Dramatic morphological changes and a burst of cell type diversification occur around post-fertilization (pf) week 3, establishing the fundamental body plan^3^. The coupled processes of neurulation and somitogenesis are central to AP axis formation^10^. By Carnegie Stage 8 (CS8, ∼18 dpf), the neural ectoderm forms a flat neural plate that folds inward to create a neural groove, flanked by emerging somites (visible from CS9). The open ends of the neural plate—known as the anterior and posterior neuropores—remain temporarily unfused. By CS10, neural tube closure initiates at multiple sites along the AP axis, including at least one at the future cervical region and another at the MHB^11^. Closure proceeds bidirectionally from these sites, with the anterior neuropore sealing by CS11 and the posterior neuropore by CS12 (∼26 dpf). At this stage, the nervous system comprises a brain, a closed spinal cord and NMPs along the AP axis, bilaterally flanked by approximately 25 pairs of somites^12^.

Recently, single cell profiling of early stage human embryos has been applied to characterize the molecular signatures of cells during the establishment of the AP axis^13–16^. However, such snapshots are unlikely to be sufficient for unraveling the underlying mechanisms, which we know to depend on a highly orchestrated set of interactions between various cell lineages and structures unfolding in space and time. For instance, the MHB functions as an organizer by producing FGF8^17^, while the somitic mesoderm synthesizes RA from retinol to establish a gradient that both patterns the hindbrain and spinal cord and promotes somite epithelialization^18^. Additional signaling pathways, such as WNT and SHH, also play well-established roles in the spatial and temporal organization of cell fates during gastrulation^19^.

While the dynamics of these and other signals have been experimentally characterized in model organisms like mouse and chick, they are challenging to study in human embryos for practical and ethical reasons. But to the extent that human data are available, they clearly suggest species-specific differences^3,19–21^, providing a strong rationale for direct studies in human materials^22^. Furthermore, the roles of direct cell-cell interactions, mechanical forces, and gene regulatory networks remain difficult to dissect in intact embryos—including model organisms but especially in humans.

The emergence of stem cell-based embryo models is unlocking new possibilities for how we study early mammalian development^3^. Mouse or human PSCs exposed to the WNT agonist CHIR form well-organized germ layers and exhibit robust axial elongation, *i.e.* gastruloids^1,23–26^. Adding other factors with specific timing, like dual SMAD inhibitors^27^, or RA and extracellular matrix (ECM; Matrigel)^4–6,28,29^, can promote co-development of the caudal neural tube and somites, but still fail to obtain anterior neural lineages. Importantly, various strategies have succeeded in producing anterior neural lineages in the context of a gastruloid or gastruloid-like model, including by introducing WNT antagonists^30^, by co-culturing with extraembryonic endoderm^31^, with microfluidic devices^32–34^, or by spatially patterning organoids’ relationships with one another^35^. Although the extended AP axis has been introduced to the mouse gastruloid through exposure to hypoxic environments^36^, to our knowledge, none of these efforts have resulted in the concurrent development of anterior neural structures alongside somitogenesis from hPSCs. As a result, a stem cell-derived model that reconstitutes advanced features of gastrulation along the full AP axis remains a major gap in our efforts to study early human embryogenesis.

Here, we leverage the apparent self-organizing potential of multiple populations of differentially pre-patterned hPSCs to obtain a new human embryo model that recapitulates key aspects of neurulation and somitogenesis, from the anterior MHB to the posterior tailbud. Specifically, we show that FGF2-pretreated hPSCs (anterior cues), when combined with (CHIR→CHIR+RA)-pretreated hPSCs (posterior cues), robustly contribute to a continuum of brain- and neural tube-like structures, conditional on the presence of the posteriorly-derived somitic mesoderm. Single cell RNA-seq (scRNA-seq) of these human “AP gastruloids” identified cell types expected in conventional^1^ or RA^4^ human gastruloids (*e.g.* somitic, cardiac and intermediate mesoderm; endoderm) but also regionalized hindbrain (*i.e.* rhombomeres) and neural crest (*i.e.* cranial, vagal, trunk). Perturbation of folic acid metabolism or rho-associated kinase (ROCK) signaling induced spinal cord defects, phenocopying aspects of spina bifida and other neural tube defects, highlighting this model’s potential for studying congenital disorders^9^. Taken together, human AP gastruloids: 1) advance our ability to model early human development and congenital disorders *in vitro*; and 2) illustrate the potential of a simple “pattern-and-mix” approach for building spatially coordinated human embyro models.

## RESULTS

### Assembly of human AP gastruloids from dual-patterned hPSC aggregates

To more fully reconstitute human AP axis patterning *in vitro*, we postulated that the exposure of hPSCs to differential morphogen cues would be critical. Current post-gastrulation embryo models—including gastruloids and trunk-like structures—rely on the homogenous activation of WNT signaling^4,5,23–26^, which restricts hPSCs to posterior fates. Thus, we hypothesized that the introduction of anterior fates to human gastruloids, *e.g.* the precursors of the brain and facial tissues, might require the modular integration of WNT-naive (anteriorized) and WNT-treated (posteriorized) hPSCs.

We first sought to identify conditions for generating anteriorized hPSCs that were capable of self-organizing with posteriorized hPSCs. To visualize the interactions of differentially pre-patterned cells, we initially used two fluorescently tagged hPSC lines (iPS11-mScarlet & iPS11-GFP). For anterior pre-patterning, iPS11-mScarlet cells were cultured in NDiff227 and exposed to various anterior cues (FGF2, XAV939 [WNT inhibition], LDN193189/SB431542 [TGFβ/BMP inhibition], no-treatment control) for 48 hrs prior to aggregation (**Extended Data Fig. 1a**). In parallel, we followed our previously reported protocol^4^ for inducing human RA gastruloids to generate posteriorized hPSCs: 24 hrs of CHIR treatment followed by 24 hrs of CHIR and RA treatment (**Extended Data Fig. 1a**). We then mixed these populations, while varying the timing of “assembly” (**Extended Data Fig. 1a**).

When anteriorized and posteriorized hPSCs were mixed at the 0-hr timepoint, *i.e.* concurrent with aggregation, they remained intermingled and disorganized throughout induction under all anteriorizing conditions except for dual TGFβ/BMP inhibition, which efficiently converts hPSC to neural cell fates^37^ (top rows of **Extended Data Fig. 1b-e**). In contrast, when anteriorized and posteriorized aggregates were assembled at either the 3-hr or 24-hr timepoint, they appeared to interact in a more organized fashion, with iPS11-mScarlet cells usually forming a brain-like domain at the anterior pole of an elongated, iPS11-GFP-derived posterior domain—regardless of the anteriorizing condition (middle & bottom rows of **Extended Data Fig. 1b-e**). However, using FGF2 as the anterior cue resulted in a more extensive interaction, with anteriorized iPS11-mScarlet cells intercalating into posteriorized iPS11-GFP to give rise to an elongated neural tube-like structure that appeared to derive from both populations. Notably, such intercalation was only observed when the assembly was performed at the 3-hr timepoint (**Extended Data Fig. 1c**). Taken together, these results demonstrate that anteriorly and posteriorly pre-treated hPSCs have the potential to extensively interact, but also that the outcome is dependent on the identity of the anterior cue and the timing of assembly.

### Live imaging of human AP gastruloid formation

To more effectively visualize the dynamics by which anteriorized and posteriorized hPSCs interact under these conditions, we switched to the RUES2-GLR cell line (SOX2::mCitrine, TBXT::mCerulean, SOX17::tdTomato)^38^ for inducing RA gastruloids^4^ for assembly with FGF2-treated iPS11-mScarlet hPSCs (**Fig. 1a**); as described further below, this switch has the added benefit of allowing us to distinguish these cell populations in scRNA-seq data based on genetic variation. FGF2-treated iPS11-mScarlet hPSCs, when assembled with RUES2-GLR-derived RA gastruloids at the 3-hr timepoint (**Fig. 1a**) gave rise to the anticipated interactions and resulting structure, *i.e.* an anterior and posterior domain, connected by an elongated neural tube with both anterior and posterior contributions (**Fig. 1b**).

**Figure 1.**
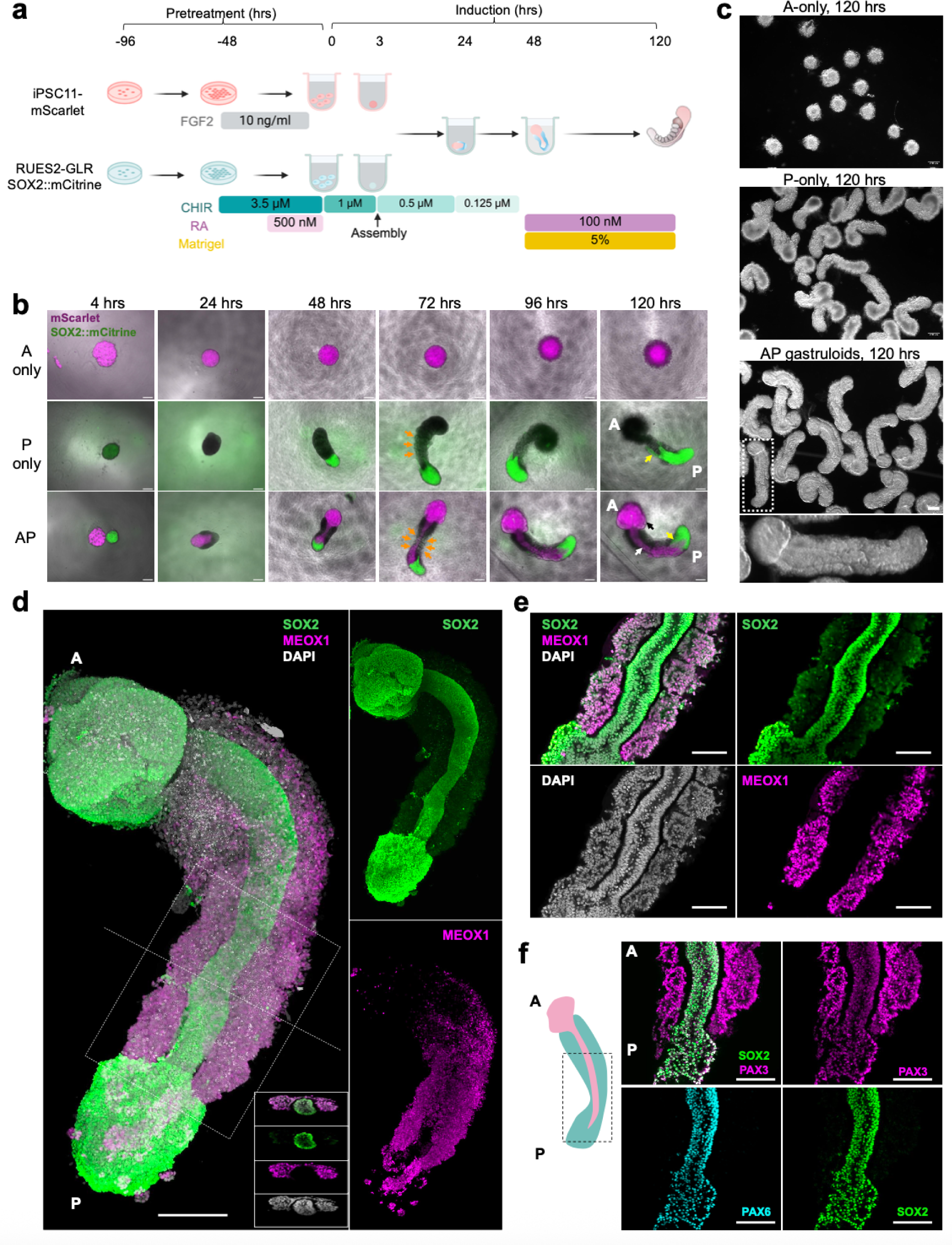
Induction of human AP gastruloids with dual patterned hPSC aggregates. **(a)** Schematic of protocol to generate AP gastruloids. After 2 days of separate pre-treatment with FGF2 or CHIR→CHIR/RA, hPSCs were dissociated into single cells and counted at 0 hrs. 2,000 anteriorized and 2,000 posteriorized hPSCs were aggregated separately in each well of a 96-well plate, followed by assembly 3 hrs later. Matrigel and additional RA were added at 48 hrs to enhance the elongation and segmentation. **(b)** Live-cell imaging of gastruloids derived from anterior (A-only, 4,000 cells/well), posterior (P-only, 4,000 cells/well), or anterior-posterior (AP) hPSC from 4 to 120 hrs. Purple corresponds to anteriorized iPS11-mScarlet cells, and green to SOX2::mCitrine from posteriorized RUES2-GLR cells. In panels corresponding to P-only and AP gastruloids at 72 hrs, orange arrows highlight segmented somites. In panels corresponding to P-only and AP gastruloids at 120 hrs, the AP axis is indicated by A and P labels, black arrows highlight neural folds in the brain-like structure, while white and yellow arrows highlight for aspects of the neural tube derived from anteriorized or posteriorized cells, respectively. Scale bar: 200 μm. **(c)** Representative brightfield images of A-only, P-only, and AP gastruloids at 120 hrs. At the bottom, we show a zoomed-in view of the AP gastruloid highlighted with a white dashed box. Scale bars: 200 μm. **(d)** Maximum projection view of a representative immunostained AP gastruloid at 120 hrs. SOX2 marks the neural lineage, and MEOX1 marks somites. A cross-section taken at the level of the white dashed line is shown in the bottom right inset of the main image (top to bottom: SOX2+MEOX1, SOX2, MEOX1, DAPI), while the top and bottom subpanels at the far right show SOX2-only and MEOX1-only views of the same gastruloid, to highlight neural tube and somite structures, respectively. Scale bar: 200 μm. **(e)** Images of a single plane of subregion of the AP gastruloid shown in panel **d** marked by a dashed white rectangle. Scale bar: 100 μm. (**f**) Images of a single plane of a subregion of a 120-hr AP gastruloid. A cartoon of that gastruloid is shown to the left with a dashed black rectangle that indicates the approximate boundaries of the view shown. SOX2/PAX6 exclusively mark the neural tube. PAX3 marks both somites and neural tube. Scale bar: 100 μm.

We sought to monitor how these subpopulations interact via live-cell imaging (**Fig. 1b**; **Extended Data Fig. 2a, Extended Data Movie.1-3**). Importantly, the formation of continuous neural tissue appeared to depend on physical contact between the anterior and posterior hPSC aggregates. In the absence of posterior cells, anterior cells formed a sphere but neither brain-like nor neural-tube-like structures (**Fig. 1b-c**). In the absence of anterior cells, posterior cells simply gave rise to RA gastruloids as previously reported^4^, with no brain-like domain (**Fig. 1b-c**). However, when anteriorized (pre-treatment: FGF2) and posteriorized (pre-treatment: CHIR→CHIR+RA) aggregates were assembled at 3 hrs, they immediately attached. At 24 hrs, the anterior cells began to ingress into the posterior aggregate, resulting in a neural-plate-like structure (**Fig. 1b**; **Extended Data Fig. 2a**). Through 48 hrs, the anterior cells continued to extend and contribute to a thin neural-tube structure, connected to the anterior brain-like structure at one pole and the posterior tailbud region at the other, suggestive of reconstitution of continuous patterning of neural tissues across the AP axis. Upon the addition of Matrigel and RA at 48 hrs, the entire structure further elongated along the AP axis, with the clear segmentation of bilateral somites and an mScarlet+ neural tube-like structure at the midline (**Fig. 1b**; **Extended Data Fig. 2a**). From 72 to 120 hrs, mScarlet+ neural-fold-like structures became visible within the anterior brain-like structure (**Fig. 1b-c**). By 120 hrs, AP gastruloids consistently included a brain-like structure at the anterior pole that was continuous with a SOX2+/PAX6+ neural tube-like structure in the trunk region, flanked bilaterally by MEOX1+/PAX3+ somites and caudally by a CDX2+ neural tail (**Fig. 1d-f**; **Extended Data Fig. 4e**).

### Morphological analysis of human AP gastruloids

We performed more extensive morphological analysis to evaluate the robustness by which the key features of AP gastruloids emerged (**Extended Data Fig. 2b-d**). By 48 hrs, symmetry breaking was observed in the vast majority of P-only (91%) and AP (95%) gastruloids (**Extended Data Fig. 2c**). By 72 hrs, elongation and somites were also evident in most P-only (71%) and nearly all AP (95%) gastruloids. Consistent with a dependency on the interaction between anteriorized and posteriorized hPSCs, brain and neural tube structures were also present in nearly all AP gastruloids at 72 hrs (95%) but in no P-only gastruloids (0%). Furthermore, as expected, none of the aforementioned structures manifested in A-only hPSC aggregates. By 120 hrs, >90% of AP gastruloids included brain-like and neural tube-like domains flanked by segmented somites (**Extended Data Fig. 2c**). These had a mean length of 1781 ± 191 μm, similar to P-only gastruloids (mean 1813 ± 180 μm) (**Extended Data Fig. 2b**).

The number of somites observed varied from 6 to 11 pairs per AP gastruloid. The first 5 somites (S1-S5) usually epithelialized within ∼40 hrs of Matrigel addition at a rate of ∼4 hrs per somite with an average size of 90 ± 16 μm (n=18 somites from n=5 AP gastruloids), recapitulating the segmentation clock in both human embryos^39^, and similar to other human gastruloids/somitogenesis models^4,25,26,21^ (**Extended Data Fig. 2d-e**). Since it has been shown to be essential in the elongation of both mouse and human posterior gastruloids^4,28^, we examined the effect of either withholding or accelerating Matrigel (**Extended Data Fig. 3a**). In the absence of Matrigel, we still observed the ingression of anteriorized hPSCs into posteriorized hPSCs, as well as the formation of a thin neural-tube-like structure (**Extended Data Fig. 3b-c**). This suggests that the neurulation from anteriorized hPSCs does not require Matrigel, in contrast with neurulation from posteriorized hPSCs^4,6^. Accelerated addition of Matrigel (24 hrs rather than 48 hrs) led to accelerated elongation (993 ± 95 μm vs. 760 ± 68 μm at 48 hrs), together with the earlier appearance of segmented somites (**Extended Data Fig. 3b-d**). However, the lengths of the resulting AP gastruloids were comparable by 120 hrs (1847 ± 185 μm vs. 1892 ± 166 μm), suggesting other factors may be limiting, *e.g.* stem cell pool size and/or morphogen diffusion^40^.

Taken together, these results show that two populations of hPSCs, when separately pre-patterned with either FGF2 and CHIR→CHIR+RA and then mixed with specific timing, robustly self-organize into a human embryo model with extended patterning along the AP axis, including a brain-like domain that is continuous with a neural tube-like domain, flanked by segmented somites.

### Neurulation dynamics in human AP gastruloids

To characterize how the neural tube-like structure observed in AP gastruloids is formed, we performed time-course immunostaining followed by CUBIC tissue clearing^41^ and confocal imaging to obtain high-resolution 3D structures (**Fig. 2a**). We stained AP gastruloids with SOX2 and MEOX1 antibodies to visualize neural and somitic lineages, respectively. As early as 24 hrs, FGF2-treated iPS11-mScarlet cells showed high expression of SOX2 and had begun to ingress into posteriorized CHIR/RA-treated RUES2-GLR cells (**Fig. 2b**). Also by 24 hrs, RUES2-GLR cells exhibited symmetry breaking, with SOX2+ cells restricted at the caudal end and a handful of MEOX1+ (a marker of early presomitic mesoderm^42^) cells adjacent to iPS11-mScarlet cells (**Fig. 2b**). These patterns were observed in all 7 AP gastruloids examined, and suggest signaling between the differentially pre-treated cell populations as they dynamically integrate. At 30 hrs, SOX2+ anteriorized cells continued to ingress into the posterior aggregate, and came in contact with posterior SOX2+ cells to form a plate-like structure (**Fig. 2c**). Also by 30 hrs, the bilaterally located MEOX1+ RUES2-GLR cells became more abundant, and a rostral to caudal gradient of increasing CDX2 expression in RUES2-GLR cells also emerged (**Fig. 2c**; **Extended Data Fig. 4a**), consistent with spatial organization of presomitic mesoderm and NMPs along the AP axis.

**Figure 2.**
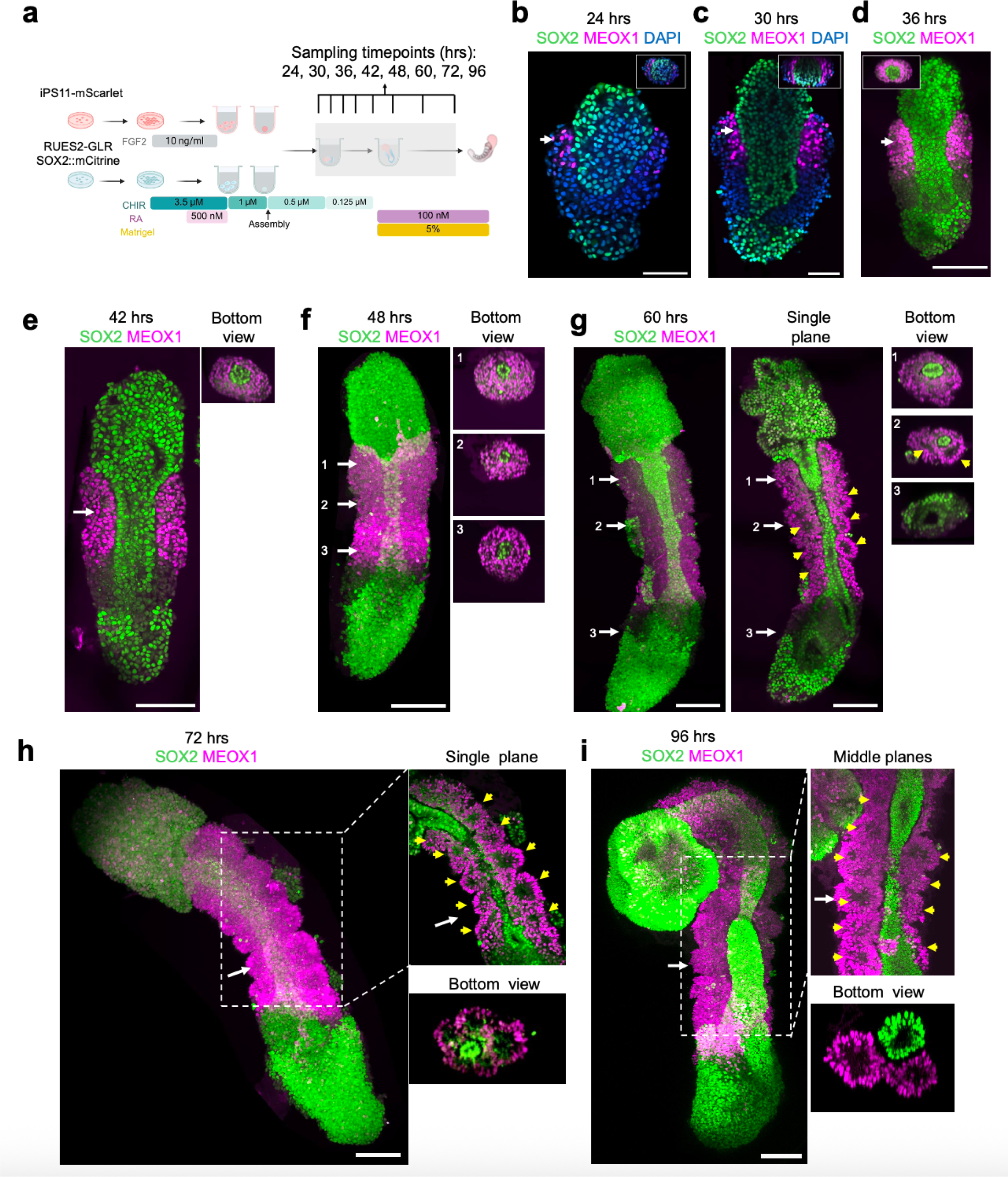
Neurulation dynamics in human AP gastruloids. **(a)** Schematic of time-course immunostaining experiment to characterize neurulation dynamics in AP gastruloids. As dramatic morphological changes were observed between 24-48 hrs in live cell imaging experiment (**Extended Data** Fig. 2a), we collected AP gastruloids every 6 hrs from 24-48 hrs, and then also at 72 and 96 hrs. **(b-i)** Representative images of AP gastruloids stained with neural (SOX2, green channel) and somitic (MEOX1, magenta channel) markers from 24 to 96 hrs. Scale bar: 100 μm. **(b-d)** Images of single planes of AP gastruloids at 24 **(b)**, 30 **(c)** and 36 **(d)** hrs. Insets at top right show cross-sections from positions of white arrows. **(e-f)** Images of AP gastruloids at 42 hrs (single plane) **(e)** or 48 hrs (maximum projection) **(f)**. At right, we show cross-sections from positions of numbered white arrows. A lumen in the neural tube was not observed at 42 hrs but became apparent in rostral (positions 1 & 2), but not caudal (position 3), cross-sections by 48 hrs. **(g)** Maximum projection (far left) and single plane (middle) images of the same stained AP gastruloid at 60 hrs. At right, we show cross-sections from positions of numbered white arrows. Yellow arrowheads highlight emerging somite segmentation in both single plane and cross-section views. A lumen in the neural tube is notably apparent in all three cross-section positions at 60 hrs. **(h-i)** Left: Maximum projection images of stained AP gastruloids at 72 **(h)** and 96 **(i)** hrs. Top right: Single plane **(h)** or maximum projection of 6 z-stacks of middle planes **(i)** of subregion (dashed white rectangle) to highlight neural tube (green) and somite (magenta) structures. Bottom right: Cross-section from position of white arrow. Yellow arrowheads highlight segmented somites.

The future neural tube domain underwent an extension and narrowing process from 30 to 48 hrs (**Fig. 2c-f**), with the formation of a lumen by 48 hrs (**Fig. 2e-f**, bottom view insets). The presence of a lumen was more apparent in the rostral half of the neural tube at 48 hrs (**Fig. 2f**, bottom view insets), but was also quite clear at the caudal end by 60 hrs (**Fig. 2g**, bottom view insets). Interestingly, in addition to the gradient of CDX2 expression in posteriorized RUES2-GLR cells, we also detected weak but evident CDX2 signals in the caudal GFP-cells from 30-48 hrs (**Extended Data Fig. 4a-c**). Furthermore, we also captured a few mScarlet+/TBXT+ cells residing in the posterior neuropore (**Extended Data Fig. 4d**), indicating a very small pool of NMPs derived from anteriorized iPS11-mScarlet cells. Of note, almost a third of the neural tube at the caudal end was CDX2+ by 120 hrs (**Extended Data Fig. 4e**), suggesting that at least by this stage, NMPs are contributing to the posterior elongation of the neural tube. However, NMPs’ contribution to the formation of the neural tube-like structure in AP gastruloids at earlier stages remains unclear.

In mouse embryos, MEOX1 is exclusively expressed in the mesoderm lineage as early as the gastrulation stage (E7.0-7.5) and becomes prominent in presomitic mesoderm and segmented somites^43^. Similarly, MEOX1 was detected in human AP gastruloids from 24 hrs and dramatically increased in expression in all mesodermal cells from 30 to 48 hrs (**Fig. 2b-f**). The segmentation of somites was only observed after the addition of the Matrigel at 48 hrs. Around 3-4 pairs of segmented somites flanking the spinal cord became visible by 60 hrs (**Fig. 2g**) and this increased to 8-11 somite pairs from 72-96 hrs with some gastruloid-to-gastruloid variation (**Fig. 2h-i**). We believe that the modestly greater number of somites observed here, as compared with our earlier morphological characterizations, is due to better imaging resolution (20x single plane immunofluorescence here [**Fig. 2h**] vs. 4x brightfield above [**Extended Data Fig. 2d-e**]). Cross-section views enabled us to quantify the sizes of somite lumens (mean 77 ± 10 μm by 72 hrs, based on n=21 somites from n=3 AP gastruloids, based on examining the z-stacks for the single plane exhibiting maximal diameter for any given lumen).

Throughout metazoans, AP axis specification is coupled to the spatiotemporal activation of HOX genes, with 3’-end genes (*e.g.* HOX1-3) activated early in the anterior region and 5’-end genes (*e.g.* HOX9-13) activated early in the posterior region^44^. To examine HOX gene expression patterns in our AP gastruloids, we dissected AP gastruloids at 120 hrs into thirds and then performed RT-qPCR on RNA derived from each section (**Extended Data Fig. 4f**). We observed a higher expression of 3’-end HOXA genes (HOXA1-3) in the anterior third of AP gastruloids, and higher expression of internal HOXA genes (HOXA4-7) in the middle third of AP gastruloids. Most 5’-end HOXA genes were not detected (HOXA9/10/13), suggesting these genes were not active yet (or a technical failure). The posterior third of AP gastruloids—predominantly NMPs—exhibited high expression of most HOXA genes. Broadly similar patterns were observed for the other HOX gene families, although none as cleanly as HOXA (**Extended Data Fig. 4f**).

Taken together, these results show that the AP axis patterning happens soon after the integration of two prepatterned hPSCs, followed by the concurrent formation of lumens in both neural tube- and somite-like structures. Interestingly, our imaging results suggest that NMPs in AP gastruloids derive from both posteriorized^4^ and anteriorized hPSCs, but with populations of both origins located at the caudal end like their *in vivo* counterparts.

### Human AP gastruloids contain the major cell lineages of neurulation and somitogenesis

To characterize cell type diversity in more depth, we profiled 8 human AP gastruloids at 120 hrs using single cell RNA sequencing (scRNA-seq; 10x Genomics). After applying stringent quality control criteria (*e.g.* UMI count >5,000, doublet removal), we obtained 12,469 single cell transcriptional profiles, with a median of 9,499 UMIs and 4,063 genes detected per cell. Based on well-established marker genes, we identified 11 main cell types, one of which remained unannotated (**Fig. 3a**, left; **Extended Data Fig. 5**; **Supplementary Table 1**). Upon iterative subclustering of brain (**Fig. 3a.I**), neural tube (**Fig. 3a.II**), neural crest (**Fig. 3a.III**) and mesodermal (**Fig. 3a.IV**) cell types, the number of annotated cell types in these human AP gastruloids expanded to 26 (**Fig. 3b**; **Supplementary Table 1**). Our confidence in these annotations was reinforced by *post hoc* integration with single cell atlases of mouse embryogenesis^7,45^ (**Extended Data Fig. 6**; **Supplementary Table 2**).

**Figure 3.**
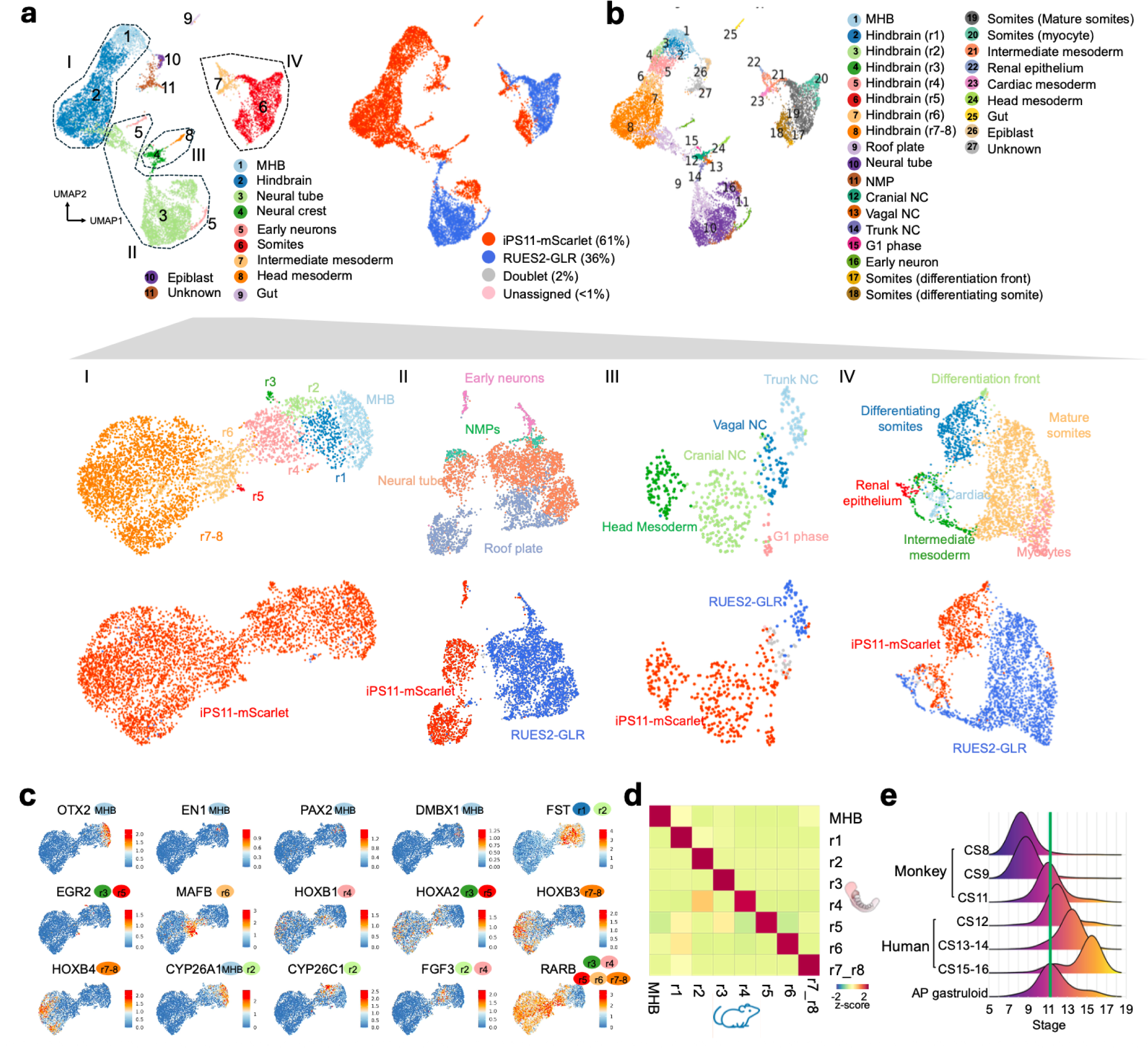
Transcriptome profiling of human AP gastruloids at single-cell resolution. **(a)** Annotation of major cell types, subtypes and cell-line-of-origin for scRNA-seq data from human AP gastruloids harvested at 120 hrs. Top left: Global UMAP of all cells passing QC (12,469 cells) numbered and colored by 11 annotation labels corresponding to major cell types. Dashed lines and Roman numerals denote four subgroups that were iteratively subclustered, resulting in subtype annotations as shown below. Top right: Identical UMAP as top left but colored by assignment to one of two hPSC lines-of-origin based on genetic variation in scRNA-seq data^60^. 61% derived from the anteriorized iPSC11-mScarlet line and 36% from the posteriorized RUES2-GLR line, while 2% were ambiguous (*i.e.* doublets) and 1% unassignable. Middle row: UMAPs of four subgroups, colored by subtype annotation, including hindbrain/MHB (I, 4,982 cells), neural tube/early neurons (II, 3,724 cells), neural crest/head mesoderm (III, 464 cells), and somitic/intermediate mesoderm (IV, 2,790 cells). MHB: midbrain−hindbrain boundary. r1-r8: Rhombomeres 1 to 8. NMP: Neuromesodermal progenitors. NC: neural crest. Bottom row: Identical UMAPs as middle row but colored by assignment to cell-line-of-origin based on genetic variation in scRNA-seq data^60^. (**b**) Identical UMAP as top left/right of panel **a** but numbered and colored by one of 27 annotation labels corresponding to cell subtypes. (**c**) Identical UMAP as subpanel I of panel **a**, showing iterative subclustering of 4,982 hindbrain/MHB cells, here colored by expression of select marker genes (raw UMI counts divided by total counts per cell, multiplied by 10,000, and natural log transformed) that were used to annotate rhomobomere subtypes. The labeled/colored circles next to each gene name indicate which cell type(s) the marker gene corresponds to, with references provided in **Supplementary Table 1**. (**d**) Annotated hindbrain/MHB subtypes were compared between E8.5 mouse embryos (*x-*axis) and 120-hr human AP gastruloids (*y-*axis) via non-negative least-squares regression. The heat map shows the combined regression coefficients (row-scaled). (**e**) For each cell from the human AP gastruloid or from each human/monkey embryonic stage (CS8, CS9, and CS11 for monkey; CS12, CS13-14, and CS15-16 for human), the 15 nearest neighbors were identified within each human or monkey embryonic stage. The distribution of stages among these 15 *in vivo* nearest neighbors is plotted. CS13–14 and CS15–16 are represented as 13.5 and 15.5, respectively, on the x-axis.

Although cell populations corresponding to forebrain and midbrain are missing, AP gastruloids contained cells resembling all other expected neural cell types spanning a *SOX2*+ continuum of the AP axis from the midbrain-hindbrain boundary (MHB) (*OTX2+, EN1+*, *WNT4+, PAX2+, DMBX1+*), hindbrain (*MAFB+*, *GBX2+*, *HOXA2+*, *HOXD4+, PAX6+*), and neural tube (*PAX6+, HOXC6+, HOXC9+*) (**Fig. 3a**, left; **Fig. 3c**; **Extended Data Fig. 5**), with the latter including both roof plate (*LMX1A+, MSX2+, WNT1+, WNT3A+*) and NMP (*CDX2+*) subtypes (**Fig. 3a.II**; **Extended Data Fig. 7a**). Additional neural derivatives, such as neural crest (*SOX10+*, *FOXD3+, ETS1+*) and early neurons (*ONECUT1+*, *ONECUT2+*) were also present (**Extended Data Fig. 5**; **Extended Data Fig. 7a**). Finally, small numbers of cells resembling gut (*SOX17+*, *FOXA2+*) and epiblast cells (*NANOG+*) were also detected (**Extended Data Fig. 5**).

Our initial annotations identified both somitic (*MEOX1+*, *TCF15+*) and intermediate (*WT1*+, *OSR1+*) mesoderm, as well as head mesoderm (*TWIST1+, FOXC2*+) (**Fig. 3a**, left; **Extended Data Fig. 5**). Iterative subclustering revealed subsets of somitic mesoderm corresponding to the differentiation front (*TBX6+, RIPPLY2+, MESP2+, MEOX2-*), differentiating somites (*PAX3+, FST+*), mature somites (*PAX3+, FST-*) and myocytes (*NEB+*), as well as subsets of intermediate mesoderm corresponding to renal epithelium (*LHX1*+, *PAX2*+, *PAX8*+) and cardiac mesoderm (*HAND2*+, *TNNT2*+) (**Fig. 3a.IV**; **Extended Data Fig. 7b**). Of note, head mesoderm appeared more closely related to neural crest than somitic/intermediate mesoderm (**Fig. 3a**, left), a topic that we return to further below.

### Human AP gastruloids contain hindbrain (rhombomere) and neural crest subtypes

Rhombomeres are compartments of the developing vertebrate hindbrain that emerge as segments along the AP axis^18,46,47^. To investigate whether these deeply conserved cellular subtypes emerge in AP gastruloids, we reanalyzed 4,982 cells corresponding to hindbrain and MHB annotations (**Fig. 3a.I**). Remarkably, we readily identified subsets expressing marker genes of all rhombomere subtypes, with differentially expressed genes including *FST* (specific to r1/2), *EGR2* (*i.e. KROX20*; specific to r3/5), *MAFB* (specific to r5/6), and various HOX genes (**Fig. 3c**). In model organisms, rhombomere segments also promote morphogen gradients along the AP axis. For example, r2/4 secrete FGF3/8 to form a secondary organization centre apart from MHB^48–53^, while anterior rhombomeres (r1-r4) express RA-degrading enzymes to form an anteriorly decreasing RA gradient^54,55,18,56^. Strikingly, these patterns also manifested among hindbrain subtypes of human AP gastruloids, including exclusive expression of *FGF3* in r2/4, strong expression of *CYP26A1* in MHB and r1/2, *CYP26C1* in r2/3, and *RARB* (RA receptor beta) in posterior rhombomeres (r4-8) (**Fig. 3c**). Furthermore, the expression patterns of the MHB and all rhombomeres annotated here are highly correlated with their counterparts in our previously reported annotations of *in vivo* mouse hindbrain segmentation at E8.5^7^ (**Fig. 3d**). Together, these results support the view that human AP gastruloids include cellular subtypes resembling the regionalized rhombomere segments of the human hindbrain.

Neural crest cells are highly migratory, multipotent progenitors that emerge from the neural tube during neurulation. NC cells are further classified as cranial, vagal, or trunk, depending on their location along the AP axis^8,57^, with these subpopulations exhibiting markedly different developmental potentials and fates. For instance, cranial neural crest is a major source of the facial mesoderm, vagal neural crest makes key contributions to the enteric nervous system and heart development, and trunk neural crest contributes to sensory and sympathetic neurons and ganglia^58,57^. To investigate whether these neural crest subtypes are present in AP gastruloids, we reanalyzed 464 single cell profiles corresponding to the neural crest and head mesoderm annotations (**Fig. 3a.III**), the former of which could be separated into cranial (*SPP1+*, *NDST3+*, *NDST4+*, *HOX-*), vagal (*HOXA3+*, *HOXB3+*, *HOXA4+*, *HOXB4+*) and trunk (*MSX1+*, *CDX2+*, *NEUROG2+*) subtypes (**Extended Data Fig. 7c**). Head mesoderm cells, which were annotated on the basis of high expression of head mesoderm markers (*TWIST1+*, *PRRX2+*, *FOXC1+*) and low expression of neural crest markers (*SOX10-*, *FOXD3*-), appeared continuous with cranial neural crest, aligning with our understanding of its origins^8^ (**Fig. 3a.III**; **Extended Data Fig. 7c**).

### *In silico* staging of human AP gastruloids through comparison to primate embryos

To assess the progression of *in vitro* derived AP gastruloids relative to *in vivo* development of primate embryos, we compared the scRNA-seq profiles generated here with published data from cynomolgus monkey (CS8–CS11)^59^ and human (CS12–CS16)^16^ embryos. Integration and co-embedding confirmed that cell type annotations were largely consistent between AP gastruloids and natural monkey and human embryos (**Extended Data Fig. 8a**). Furthermore, by checking the developmental stage of the *in vivo* nearest neighbors of *in vitro* derived cells, we found that 120-hr human AP gastruloids were most closely aligned with the CS11-CS12 stage of primate development (**Fig. 4e**). Some cell types exhibited heterogeneity in their aligned developmental stages. For example, while neural tube and neural crest cells were strongly enriched at CS11–CS12, MHB and subsets of rhombomeres exhibited a much broader range of aligned stages, while epiblast and gut cells tended to align with earlier stages (**Extended Data Fig. 8b**). Of note, our scRNA-seq profiles derive from a batch of eight 120-hr AP gastruloids, such that this may reflect inter-gastruloid heterogeneity, intra-gastruloid heterogeneity or some combination thereof.

**Figure 4.**
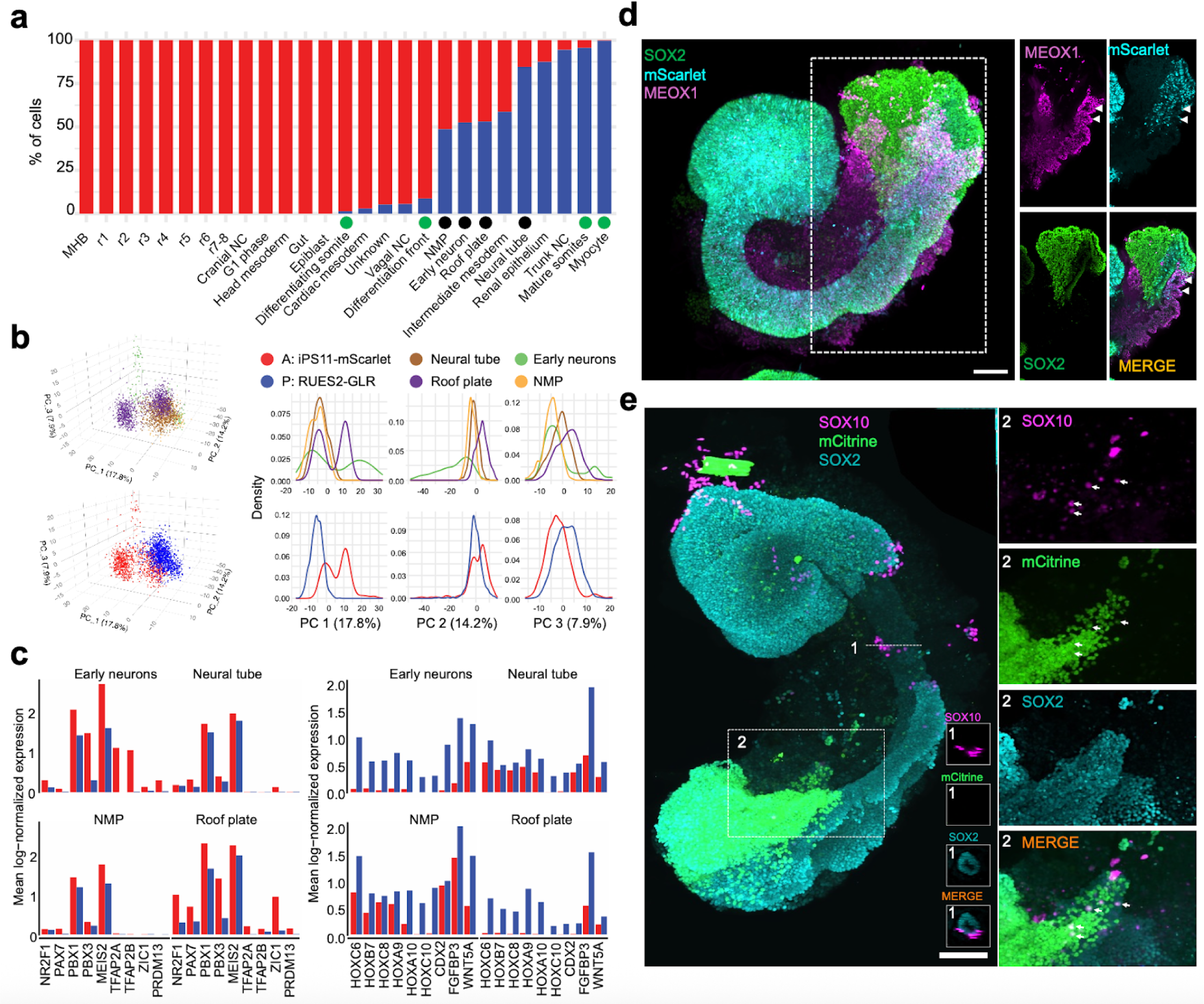
Genetic disambiguation of human AP gastruloids. **(a)** A total of 12,194 cells (97% of all cells) were assigned to one of the two hPSC lines of origin using genetic variation detected in scRNA-seq data^60^. For each of the 27 annotated cell types, the proportion of cells confidently assigned to iPS11-mScarlet (anteriorized) or RUES2-GLR (posteriorized) is displayed. Four neural tube-related cell types are marked with black dots, and four paraxial lineage cell types are marked with green dots. **(b)** Left: 3D visualization of the top three principal components (PCs) of gene expression variation in cells from four neural tube-related cell types (neural tube, early neurons, roof plate, and NMPs) calculated on the basis of the 2,500 most highly variable genes. Cells are colored by cell type (top) or by their assigned hPSC line of origin (bottom). Right: Density plots of the top three PCs, stratified by cell type (top) or by hPSC line of origin (bottom). **(c)** Averaged log-normalized expression of selected genes in four neural tube-related cell types was plotted separately for cells assigned to iPS11-mScarlet vs. RUES2-GLR. Gene expression values were calculated from original UMI counts normalized to total UMIs per cell, followed by natural-log transformation. **(d)** Imaging of a representative AP gastruloid at 120 hrs with endogenous mScarlet signal (from iPS11-mScarlet cells), stained somitic (MEOX1) and neural (SOX2) markers. Left: maximum projection of merged channels. Right: Enlarged single plane of a subregion (white dashed rectangle on the left) in separated (MEOX1, mScarlet, SOX2) and merged (MERGE) channels as indicated. White arrows highlight MEOX1+/mScarlet+ somites at the caudal end, suggesting that iPS11-mScarlet cells mainly contributed to late (posterior) somites, while RUES2-GLR cells mainly contributed to early (anterior) somites. Scale bar: 100 μm. **(e)** Imaging of an AP gastruloid at 120 hrs with endogenous mCitrine signal (SOX2:mCitrine from RUES2-GLR cells), stained neural crest (SOX10) and neural (SOX2) markers. SOX10+ neural crest cells distributed along the AP-axis and mCitrine+ RUES2_GLR cells mostly located at the caudal end. Left: maximum projection of the whole AP gastruloid. Right bottom corner in the left panel: cross-sections at the position 1 (indicated as dash line) with separated (SOX10, mCitrine, SOX2) and merged (MERGE) channels to highlight the delamination of neural crest cells from the neural tube. Right: Projection of substacks in the zoomed region 2 (indicated as dashed rectangle) with separated (SOX10, mCitrine, SOX2) and merged channels. White arrows stand for SOX10+/mCitrine+ cells. Scale bar: 100 μm.

Taken together, our analyses presented thus far demonstrate that human AP gastruloids contain cell types from all three canonical germ layers in a single structure, as well as neural crest. These cell types are well aligned with their equivalents in natural embryos from mouse, monkey and human, and at least based on their transcriptional states, cells within 120-hr AP gastruloids are most similar to CS11-CS12 primate embryos. The neural cell types span a continuum of the AP axis from the MHB to the neural tube, and several major classes of mesodermal lineages are present as well (*i.e.* head, intermediate, paraxial). We also observe AP axis defined subtypes of both hindbrain (*i.e.* rhombomeres 1-8) and neural crest (cranial vs. vagal vs. trunk), as well as evidence for further differentiation of many lineages towards more mature cell types, *e.g.* somitogenesis (*PAX3*+ somites), myogenesis (*NEB*+ skeletal myocytes; *TNNT2*+ cardiomyocytes), nephrogenesis (*PAX2/8+* renal epithelium) and neurogenesis (*ONECUT1/2+* early neurons).

### Genetic disambiguation of the contributions of anteriorized vs. posteriorized hPSCs

To investigate whether each cell type observed in 120-hr human AP gastruloids derived from anteriorized hPSCs, posteriorized hPSCs, or some mixture thereof, we leveraged genetic differences between the iPSC11 and RUES2 cell lines^60^. We confidently assigned 97% of profiled cells, including 61% to iPS11-mScarlet (anteriorized) and 36% to RUES2-GLR (posteriorized) cells (**Fig. 3a**, right; **Fig. 4a**). As AP gastruloids were initiated from pre-patterned cells at a 50/50 ratio (counted at 0 hours, mixed at 3 hours), which could be due to slightly faster expansion of the anteriorized hPSCs than the posteriorized hPSCs over the ensuing 5 days, or alternatively to technical biases during dissociation or other steps (**Fig. 1a**). Interestingly, the anteriormost cell types, (including MHB, hindbrain, head mesoderm) derived purely from anteriorized iPSC11-mScarlet cells, while most other cell types (including NMPs, paraxial/intermediate/cardiac mesoderm, neural tube) derived from <u>both</u> anteriorized and posteriorized hPSCs, but in highly varying proportions (**Fig. 4a**).

The cell line contributions were most balanced in four neural tube-related cell types (black dots in **Fig. 4a**). However, these were clearly separated in UMAP visualizations (**Fig. 3a.II**), which could be due to: (i) cell line differences, *i.e.* iPSC11 vs. RUES2; (ii) spatial heterogeneity, *i.e.* anterior vs. posterior differences; or (iii) artifacts of UMAP visualization. To evaluate this more carefully, we extracted these cell types (roof plate, neural tube, NMPs, early neurons) and performed principal components analysis (PCA) (**Fig. 4b**). The first PC (PC1: 18% of total variance) separated cells into two populations, one composed solely of iPSC11-mScarlet-derived roof plate and neurons (high PC1), and the other derived from both cell lines and all four cell types (low PC1). PC1 positively correlated genes were enriched for TFs relevant to anterior neural tube development (*e.g. NR2F1, PAX7, PBX1, PBX3, MEIS2*)^62–64^, while PC1 negatively correlated genes were enriched for TFs relevant to posterior neural tube development (*e.g. CDX2*, various HOX genes, *WNT5a*)^65–67^ (**Fig. 4c**; **Supplementary Table 3**). Taken together, these patterns suggest that the “high PC1” cluster corresponds to the anterior aspect of the neural tube, and derive solely from anteriorized hPSCs, while the “low PC1” cluster corresponds to the posterior aspect of the neural tube, and derives from both anteriorized and posteriorized hPSCs. This interpretation is consistent with our live-cell imaging of AP gastruloid formation, in which anteriorized hPSCs ingress into posteriorized hPSCs during the formation of the neural tube-like structure, rather than the other way around (**Fig. 1b**; **Extended Data Fig. 2a**).

Considered together, the paraxial lineages in 120-hr AP gastruloids also derived from both anteriorized and posteriorized hPSCs, but in contrast with the neural tube lineages, each cell type overwhelmingly arose from one or the other (green dots in **Fig. 4a**). Specifically, early somitic cell types (differentiation front, differentiating somites) derived from anteriorized hPSCs (>90%), while mature somitic cell types (mature somites, mycotes) derived from posteriorized hPSCs (>90%) (**Fig. 3a. IV**; **Fig. 4a**). A parsimonious interpretation is that early somites derive from posteriorized hPSCs (and thus are mature by 120 hrs) and late somites from anteriorized hPSCs (and thus are still differentiating at 120 hrs). As new somites are expected to arise caudally and move anteriorly as they differentiate, this is also consistent with our live-cell imaging, in which somites were mScarlet-negative at 72 hrs, but both mScarlet-negative (anterior, mature) and mScarlet-positive (posterior, still differentiating) at 120 hrs (bottom row of **Fig. 1b**). Further consistent with this interpretation, staining of 120-hr AP gastruloids found that MEOX1+/mScarlet+ cells were exclusively caudal (**Fig. 4d**). Interestingly, the intermediate mesoderm lineage exhibits a similar pattern to paraxial mesoderm in that although both cell types contribute, the more differentiated cells (*i.e.* renal epithelium) were more likely to originate from posteriorized hPSCs (**Fig. 3a. IV**; **Fig. 4a**).

Like paraxial mesoderm, the neural crest lineage of 120-hr AP gastruloids derived from both prepatterned cell lines, but individual cell types overwhelmingly arose from one or the other. Specifically, nearly all assignable cranial neural crest (100%), head mesoderm (100%), and vagal neural crest (94%) cells derived from anteriorized iPS11-mScarlet hPSCs, while nearly all trunk neural crest (94%) derived from posteriorized RUES2-GLR hPSCs (**Fig. 3a.III**; **Fig. 4a**). To check this visually, we stained AP gastruloids with both pan-neural (SOX2) and neural crest (SOX10) markers and imaged them together with the endogenous SOX2:mCitrine from RUES2-GLR line. SOX10+ neural crest cells were associated with brain- and neural tube-like structures throughout the AP axis, but SOX10+ cells within the caudal region were enriched for also being mCitrine+ (**Fig. 4e**), although mCitrine signal was heterogeneous in the rostral region across individual AP gastruloids. Taken together, these results showcase the general consistency of neural crest subtype annotations (based on gene expression), lineage annotations (based on genotype) and physical localization (based on imaging), along the AP axis.

### Comparison of human AP gastruloids vs. human RA gastruloids

We next sought to compare the cellular composition of AP vs. RA gastruloids^4^. As a context refresher, human RA gastruloids are a system that we recently described, which along with several other recently described systems are capable of concurrent modeling of somite and neural tube structures^4–6^. In most of our reported experiments, RA gastruloids were derived from the same cell line (RUSE2-GLR) and a similar protocol as was used here for posteriorized hPSCs (**Extended Data Fig. 9a**). Thus, we reasoned that a comparison of new and published data might shed light on whether and how anteriorized hPSCs shift the cell fate distribution of posteriorized hPSCs in AP gastruloids.

We co-embedded our scRNA-seq data from 120-hr AP gastruloids and 120-hr RA gastruloids^4^ (**Extended Data Fig. 9b-c**), while also examining the cell fate distribution of the RUES2-GLR line in both models (**Extended Data Fig. 9d**). As expected, epiblast and anterior cell lineages (*e.g.* brain-like cells, cranial neural crest, head mesoderm) were only detected in AP gastruloids, consistent with the fact that they originate solely from anteriorized hPSCs (**Fig. 4a**; **Extended Data Fig. 9b-c**). In contrast, endothelium was the only cell type detected in RA gastruloids but not AP gastruloids.

All other lineages were detected in both models and co-clustered in the co-embedding. However, a few key differences are worth noting. First, endoderm and cardiac mesoderm were overwhelmingly derived from anteriorized hPSCs in AP gastruloids (100% and 97%, respectively), and the latter was much less abundant and much less mature than the cardiac mesoderm lineage in RA gastruloids (**Extended Data Fig. 9b-d**). Second, as discussed in the previous section, for both paraxial and intermediate mesoderm lineages in AP gastruloids, less mature states tended to be derived from anteriorized hPSCs, and more mature states from posteriorized hPSCs (**Fig. 3a. IV**; **Fig. 4a**). However, in RA gastruloids, the continuum of these lineages derived from the only source present, *i.e.* posteriorized RUSE2-GLR cells (**Extended Data Fig. 9d**).

Taken together, these analyses confirm the greater complexity of cell lineages, types, and states in AP gastruloids as compared with RA gastruloids^4^. Furthermore, they suggest that the presence of anteriorized hPSCs may restrict or impede the differentiation of posteriorized hPSCs in certain lineages, but without compromising the diversity of cell types present in the resulting structures.

### Modeling neural tube defects through chemical perturbation of human AP gastruloids

Neural tube defects (NTDs) are among the most common developmental disorders, collectively impacting around 1 in 1,000 newborns and causing 140,000 new cases annually^68^. It is very well established that folic acid supplementation, before and during early pregnancy, substantially reduces the risk of NTDs^69–74^, and that folic acid antagonists can cause severe NTDs, as well as congenital malformations of the heart and limbs^75,76^. Although folic acid’s role in nucleotide synthesis is deeply understood^77^, a mechanistic understanding of how modulations of this biochemistry underlie the causation or mitigation of NTDs, a developmental phenotype, remains elusive. This may be in part because of the lack of a tractable human model system for studying NTDs. Towards addressing this gap, we sought to evaluate whether human AP gastruloids could serve to recapitulate NTD dynamics *in vitro*.

First, we added aminopterin, a folic acid antagonist and antineoplastic drug that causes NTDs^75,76^, into the media during AP gastruloid formation (24 to 120 hrs), and performed immunostaining of neural (SOX2) and somitic (MEOX1) markers at 72 and 120 hrs (**Fig. 5a**). At 72 hrs, although a neural tube structure with both a lumen and segmented somites was evident, the neural tube was much thinner in the presence of aminopterin and some break points were observed (**Fig. 5b-c**). By 120 hrs, striking phenotypes were evident (**Fig. 5e-f**). Most notably, while a tube-like structure remained present immediately caudal to the anterior brain-like region, the middle section of the neural tube was missing in all 8 aminopterin-treated AP gastruloids, leaving two disconnected SOX2+ cell populations at the ends (**Fig. 5f,h**; **Extended Data Fig. 10b**). Furthermore, the anterior brain-like region was markedly reduced (41% loss by area) and the neural folds observed in controls were missing (**Extended Data Fig. 10a-b**; **Extended Data Fig. 11a-b**). Although somite segmentation remained robust, somites were smaller (13% loss by diameter; **Extended Data Fig. 11d**) and the overall length of aminopterin-treated AP gastruloids was reduced (21% shorter; **Extended Data Fig. 11c**). Together, these results show aminopterin reduces the size of AP gastruloids without strictly preventing neural tube lumen formation or somite segmentation, consistent with a model in which impaired DNA synthesis inhibits cell proliferation, but in a way that the neural lineages may be most sensitive to.

**Figure 5.**
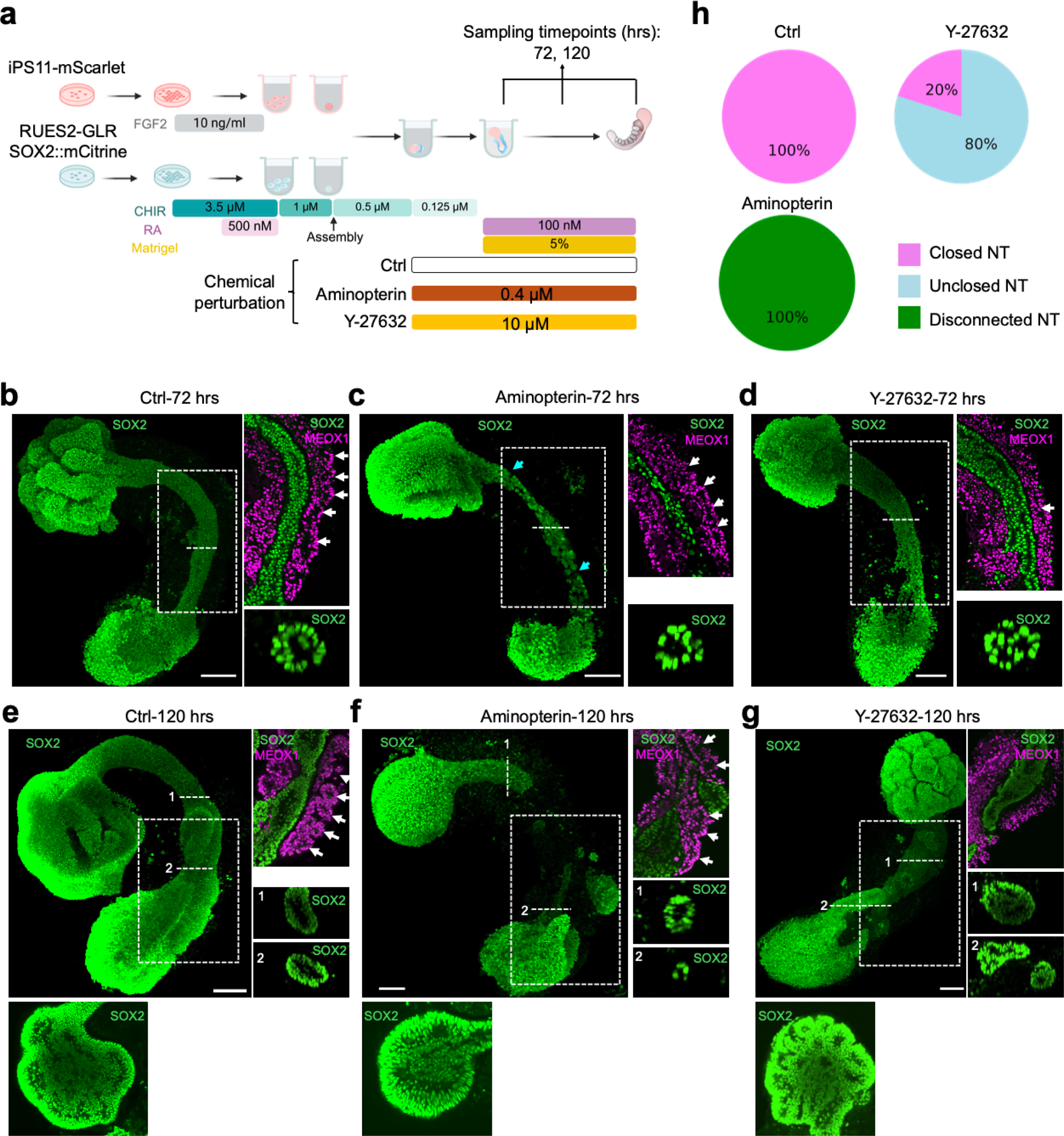
Modeling neural tube defects through chemical perturbation of human AP gastruloids. **(a)** Schematic of experiments. A folic acid antagonist (aminopterin, 0.4μM) or ROCK inhibitor (Y-27632, 10μM) was added to the induction media from 24 to 120 hrs of the AP gastruloid protocol, or no chemical treatment as a negative control (Ctrl). For characterization, 8 to 12 AP gastruloids from each group were sampled at 72 and 120 hrs, and co-stained with markers for pan-neural lineage (SOX2) and somitic mesoderm (MEOX1). **(b-d)** Representative images of AP gastruloids from each condition (indicated on the top) at 72 hrs stained for somitic (MEOX1, magenta) and neural (SOX2, green) markers. Left: maximum projection of SOX2 channel. In panel **c**, cyan arrows highlight emerging break points. Right top: Enlarged single plane of a subregion (white dashed rectangle in the left panel) in merged channels of MEOX1 and SOX2. White arrows highlight segmented somites. Right bottom: cross-sections of SOX2 channel from positions of dashed white lines in left panel for better visualization of neural tube lumen. Scale bar: 100 μm. **(e-g)** Representative images of AP gastruloids from each condition (indicated on the top) at 120 hrs stained for somitic (MEOX1, magenta) and neural (SOX2, green) markers. Left top: maximum projection of SOX2 channel. Left bottom: a middle plane of the brain-like structure to highlight neural folds. Right top: Enlarged single plane of a subregion (white dashed rectangle in the left panel) in merged channels of MEOX1 and SOX2. White arrows highlight segmented somites. Right bottom: cross-sections of SOX2 channel from positions of numbered dashed white lines in top left panel for better visualization of neural tube lumen. Scale bar: 100 μm. (**h**) Quantified categories of neural tubes in each condition. Closed NT: neural tube with lumen and without broken branches. Unclosed NT: neural tube without lumen or containing broken branches or both. Disconnected NT: neural tube missing the middle section.

Next, we examined the effect of Y-27632, an inhibitor of Rho-associated protein kinase (ROCK), on human AP gastruloid formation (**Fig. 5a**). *In vivo*, Rho GTPases bind and activate ROCK for cytoskeletal remodeling, and the genetic ablation of Rho GTPases, such as Rac1 or Cdc42, results in failure of cell protrusion and fusion^78^. Pharmacological inhibition of ROCK using Y-27632 causes NTDs in mouse embryos^9^ and disrupts somitogenesis in chick embryos^79^. We added Y-27632 during AP gastruloid formation (24 to 120 hrs), and once again performed immunostaining of neural (SOX2) and somitic (MEOX1) markers at 72 and 120 hrs (**Fig. 5a**). Consistent with observations in mouse embryos, ROCK inhibition compromised lumen formation within the neural tube structure at both 72 and 120 hrs (**Fig. 5d,g**). In contrast with aminopterin, the anterior brain-like region remained unchanged in size and neural folds remained present, albeit more numerous and smaller than in controls (**Fig. 5e,g**; **Extended Data Fig. 10a,c**; **Extended Data Fig. 11a-b**). At the caudal end of 8 of 10 Y-27632-treated AP gastruloids, multiple neural tube branches were evident, mostly disconnected such that they left an open area in the caudal region (**Fig. 5g**; **Extended Data Fig. 10c**). Overall, only 3 of 10 Y-27632-treated AP gastruloids bore a neural tube lumen structure, and 1 of these 3 had broken branches (**Extended Data Fig. 10c**), such that the rate of successful neural tube closure was decreased to 20% (**Fig. 5h**). As with aminopterin treatment, the overall length of AP gastruloids was reduced by Y-27632 (12% shorter; **Extended Data Fig. 11c**). However, somite segmentation was specifically ablated by Y-27632 (**Fig. 5d,g**). A key point is that SOX2 and MEOX1 expression levels were not altered by Y-27632 (**Fig. 5d,g**), indicating that Y-27632 does not disrupt lineage commitment, and presumably achieves its phenotypic effects by inhibiting other processes, such as cell movement^80^, in both neural and somitic lineages.

Although a comprehensive mechanistic exploration is beyond the scope of this manuscript, these results demonstrate that human AP gastruloids respond distinctly to folic acid antagonists and ROCK inhibitors (**Fig. 5h**). Our data are consistent with the hypotheses that aminopterin primarily affects cell proliferation while Y-27632 primarily impairs cell movement and morphogenetic processes. Furthermore, our findings suggest that AP gastruloids can effectively model NTDs, recapitulating key features of these disorders observed *in vivo* in the presence of clinically relevant perturbations, including aminopterin exposure.

## DISCUSSION

Early human embryonic development during neurulation and somitogenesis remains largely inaccessible for direct study due to practical and ethical constraints, while even model organisms such as the mouse present significant technical challenges at these critical stages. Recently, hPSC-derived models such as gastruloids^1^ have emerged as essential alternatives for investigating early human development. However, although gastruloids^1^ and related models^4–6,24–26,35^ reconstitute many aspects of germ layer emergence and coordination, they lack advanced anterior neural structures—a limitation likely attributable to WNT agonist exposure during their generation. This absence prevents comprehensive modeling of AP axis patterning, which is particularly problematic for studying developmental disorders affecting the full neural axis, *e.g.* anencephaly, cerebellar malformations, and spina bifida. As such NTDs represent major causes of morbidity and mortality, there remains a critical need for improved hPSC-derived models that recapitulate coordinated AP axis development. Here, we report a simple yet robust “pattern-and-mix” method for deriving gastruloids that capture a broad spectrum of AP axis features. Human AP gastruloids are generated from two populations of hPSCs—one pre-patterned with anterior cues (FGF2) and the other with posterior cues (CHIR and RA)—that, when combined with appropriate timing, robustly self-organize into continuous, elongated 3D structures (**Fig. 1d**).

Human AP gastruloids contain diverse cell types from all three germ layers—neural (including MHB, hindbrain, neural tube, neural crest, early neuron), mesodermal (including head, somitic, cardiac, intermediate), and endodermal (gut) lineages (**Fig. 3b**)—with the majority of cell types aligning most closely with the CS11-CS12 stage of primate development (**Fig. 4e**). The cellular complexity of AP axis patterning achieved greatly exceeds our previously reported human RA gastruloid model^4^, as evidenced by the detection of regionalized hindbrain subtypes (rhombomeres 1–8), regionalized neural crest subtypes (cranial, vagal, trunk), and a distinct head mesoderm subpopulation. Remarkably, these subtypes exhibit expression patterns of morphogen gradients and organizing centers that mirror their *in vivo* counterparts, *e.g.* exclusive *FGF3* expression in rhombomeres 2/4 and RA-degrading enzyme expression in anterior rhombomeres. This level of regional specification demonstrates that the pattern-and-mix strategy successfully reconstitutes not only cellular diversity but also at least some aspects of the molecular mechanisms underlying axis patterning. Furthermore, genetic demultiplexing reveals that while the anteriormost cell types (MHB, hindbrain, head mesoderm) derive solely from anteriorized hPSCs, other lineages show substantial contributions from both progenitor populations.

A key advantage of the pattern-and-mix strategy is its combination of simplicity, robustness, and scalability. By co-aggregating two pre-patterned hPSC populations, we can generate embryonic-like structures with over 90% efficiency (**Fig. 1c**; **Extended Data Fig. 2c**), and with no need for specialized equipment such as micropatterned plates^35,81^ or microfluidic devices^32–34^. This operational simplicity represents a significant advance over current methods that depend on precise geometric constraints or complex fluid handling systems to achieve AP patterning, and makes it accessible to more labs worldwide. These attributes of the protocol also make it amenable to large-scale genetic and chemical screens, especially those directed at understanding developmental mechanisms or modeling NTDs. As a proof-of-concept, exposing AP gastruloids to inhibitors of folic acid metabolism (aminopterin) and ROCK signaling (Y-27632) induced distinct neural tube defect phenotypes (**Fig. 5h**) that recapitulate the effects of these compounds observed *in vivo*.

Building on this proof-of-concept, we envision further scaling of chemical and genetic perturbations of human AP gastruloids to systematically dissect the molecular pathways governing human neurulation and somitogenesis. Beyond high-throughput applications, we note the pattern-and-mix strategy provides a potentially powerful framework for dissecting developmental mechanisms by introducing separate genetic perturbations into anterior vs. posterior progenitors to create “structured genetic mosaics”. This approach would enable targeted studies of region-specific gene function while bypassing the complex genetic crosses required for conditional knockout models in animals^82^. This same principle underpinned our tracking of the regional contributions of progenitors to specific lineages, providing an entry point for investigating how cell-cell interactions between these populations mediate their integration into cohesive structures such as a continuous neural tube. Taken together, these capabilities and possibilities position AP gastruloids as a versatile platform for systematically dissecting human embryogenesis and modeling developmental disorders.

An intriguing aspect to consider is the mechanisms underlying neural tube elongation in AP gastruloids. Time-lapse imaging shows that upon fusion of anteriorized and posteriorized hPSCs, anteriorized cells intercalate into posteriorized cells, and these populations together undergo elongation and narrowing to form a neural tube continuum flanked by segmented somites (**Fig. 2**). This neural tube extension appears to rely on convergent extension (CE)^83^, a fundamental morphogenetic process with two distinct cellular modes: a ROCK-dependent “contraction mode” involving actomyosin contractility, and a ROCK-independent “crawling mode” driven by actin-rich protrusions that allow cells to actively pull themselves between their neighbors^84,85^. Notably, AP gastruloids maintained neural tube elongation even under ROCK inhibition (**Extended Data Fig. 11c**), suggesting that ROCK-independent mechanisms may predominate in this context. Additionally, physical forces from surrounding paraxial mesoderm may contribute through compressive “squeezing”^86^, as SOX2+ neural cells derived from anteriorized hPSCs only elongate and narrow when surrounded by posterior hPSC-derived mesoderm (**Fig. 2**). These observations position AP gastruloids as a useful platform for investigating the mechanical and cellular mechanisms underlying human neural tube formation, including interactions with surrounding, non-neural tissues during extension.

AP gastruloids have important limitations that should be acknowledged. Most notably, they lack forebrain and midbrain structures, as well as a complete dorsal-ventral axis, including the notochord and ventral cell types such as floor plate neurons. The absence of these structures limits the model’s ability to fully recapitulate the 3D organization of the developing human embryo and certain signaling pathways, such as SHH-mediated ventral patterning. Additionally, while AP gastruloids capture many aspects of early human development, they represent a simplified system that may not fully reflect the complexity of tissue interactions and mechanical forces present in intact embryos. Nevertheless, these limitations also point toward promising future possibilities. In particular, recent advances in generating forebrain and midbrain progenitors^87^ and notochord-like structures^88,89^ suggest that further extensions of the pattern-and-mix strategy could potentially address these gaps. For instance, incorporating anteriorly patterned cells directed toward forebrain fates, or including ventrally patterned populations that give rise to notochord, could enhance the model’s completeness.

In summary, human AP gastruloids demonstrate that differentially patterned hPSC populations can self-organize into coordinated embryonic structures with extended AP patterning. The simplicity and scalability of this pattern-and-mix approach, combined with its amenability to genetic and chemical perturbations, establish AP gastruloids as a valuable platform for investigating human embryogenesis and modeling developmental disorders. Notably, the successful recapitulation of NTD phenotypes through chemical perturbation demonstrates the model’s potential for studying congenital malformations. More broadly, the pattern-and-mix strategy provides a generalizable framework that could be extended to other developmental contexts by incorporating different patterned cell populations. While limitations remain, this work provides a foundation for understanding fundamental mechanisms of human body axis formation and offers new opportunities for studying developmental disorders affecting early human development.

## Supplementary Tables

Supplementary Table 1. Gene markers used to annotate each of the 27 cell types in the AP gastruloids.

Supplementary Table 2. The percentages of nearest neighbor cells across cell types from natural mouse embryos are shown for each AP gastruloid cell type, listing only the five most abundant cell types.

Supplementary Table 3. Genes showing significant correlation with PC1 in the combined set of four neural tube related cell types.

Supplementary Table 4. List of antibodies. Supplementary Table 5. Oligonucleotide sequences.

## Data & code availability

The data generated in this study can be downloaded in raw and processed forms from the NCBI Gene Expression Omnibus under accession number <u>GSE308289</u>. The code used here is available at: https://github.com/shendurelab/AP_human_gastruloids.

## Supporting information

Supplementary Table 1-References of lineage-specific marker genes

Extended Data Movie 1-Live cell imaging of AP gastruloids induction-A+P

Extended Data Movie 2-Live cell imaging of gastruloids induction-P-only

Extended Data Movie 3-Live cell imaging of gastruloids induction-A-only

## Acknowledgments

We thank the members of the Shendure and Hamazaki labs for their helpful discussions. This work was supported by the Washington Research Foundation (postdoctoral fellowship to Z.L.), the Brotman Baty Institute for Precision Medicine, and grants from the Allen Family Philanthropies (Allen Discovery Center for Cell Lineage Tracing to J.S.), the Seattle Hub for Synthetic Biology, a collaboration between the Allen Institute, Chan Zuckerberg Initiative and University of Washington (award number CZIF2023-008738 to J.S.), Alex’s Lemonade Stand Foundation (Grant 19-15730), Crazy 8 Initiative (to J.S.), National Institutes of Health (1UM1HG011586 to J.S.; R01HG010632 to J.S.), and The Royalty Research Fund (to N.H.). J.S. is an Investigator of the Howard Hughes Medical Institute. We disclose that language editing, proofreading and coding were supported by AI-based tools; these were not used for conceptual development or primary manuscript writing. The authors take full responsibility for the contents of this manuscript.

## Author Contributions

Z.L., J.S., N.H. designed the research.

Z.L. performed experiments with assistance from C.K., S.X., R.M.D., E.K.N., C.L.

Z.L., C.Q. performed computational analyses with assistance from C.K., W.Y., A.V., M.B.O.

Z.L., C.Q., J.S., N.H. collaboratively explored and annotated the data.

Z.L., C.Q., J.S., N.H. wrote the manuscript.

J.S., N.H. supervised the project.

## METHODS

### Ethics Statement

Our research on the induction and molecular analysis of human RA and AP gastruloids was reviewed and approved by the Embryonic Stem Cell Research Oversight of the University of Washington (E0047-001) and is in compliance with the principles laid out by the International Society for Stem Cell Research Guidelines for Stem Cell-based Embryo Models^90^. No new experiments involving human embryos were performed in this study, and all natural human embryo data analysed were obtained from publicly available datasets.

### Statistical analyses

For box plots in this study, the thick horizontal line, upper and lower box edges, whisker and dots beyond the whisker represent the median, third quartiles (Q3), first quartiles (Q1), largest to smallest values within 1.5 times the interquartile range, and outliers, respectively. Student’s t-test was used for statistical tests. The number of replicates (N) is listed in the legends of each figure. No experimental data were excluded from the analyses.

### Human pluripotent stem cell (hPSC) culture

The RUES2-GLR human ESC line was kindly provided by A. Brivanlou (Rockefeller University). For the maintenance, 12-well plates were pre-coated with DPBS (Thermo Fisher, 14190144) containing 1% Geltrex (120 to 180 μg/ml, Thermo Fisher, A1413302) for at least 15 minutes (mins). RUES2-GLR cells were dissociated by StemPro Accutase (Thermo Fisher, A1110501) at 37 °C for 4 mins, counted, and seeded at 100,000 cells per well in StemFlex™ Medium (Thermo Fisher, A3349401) supplemented with 10 μM Y-27632 (Sellek, S1049). Cells were cultured in a 37 °C, 5% CO_2_ incubator. The medium was replaced by fresh StemFlex™ Medium without Y-27632 from the second day, and cells were passaged every three days.

The iPS11-mScarlet and iPS11-eGFP human iPSC lines were kindly provided by Dr. Michael Garton at the University of Toronto^91^. For the maintenance, 12-well plates were pre-coated with DPBS containing 5 μg/ml Vitronectin (Thermo Fisher, A31804) for at least 1 hour. The iPS11-mScarlet and iPS11-eGFP cells were dissociated by StemPro Accutase (Thermo Fisher, A1110501) at 37 °C for 4 mins, counted, and seeded at 100,000 cells per well in Essential 8™ Flex Medium (Thermo Fisher, A2858501) supplemented with 10 μM Y-27632 (Sellek, S1049). Cells were cultured in a 37 °C, 5% CO_2_ incubator. The medium was replaced by fresh Essential 8™ Flex Medium without Y-27632 daily from the second day, and cells were passaged every three days.

### Induction of human AP gastruloids

#### 1. Induction of anteriorized and posteriorized progenitor cells

For the induction of AP gastruloids, RUES2-GLR and iPS11-mScarlet cells were separately pre-treated in two distinct conditions. On day 0, RUES2-GLR and iPS11-mScarlet cells were dissociated by StemPro Accutase and seeded into Vitronectin-coated 12-well plates at a density of 15,000 and 20,000 cells per well, respectively, in NutriStem® hPSC XF Medium (Biological Industries, 05-100-1A) supplemented with 10 μM Y-27632. On day 1, medium was replaced with fresh NutriStem® medium supplemented with 5 μM Y-27632. On day 2, the medium was replaced with fresh NutriStem® medium containing 3.5 μM CHIR (Millipore, SML1046) for RUES2-GLR (posterior) and NDiff 227 medium (Takara, Y40002) containing 10 ng/ml bFGF (Thermo Fisher, PHG0360) for iPS11-mScarlet (anterior). On day 3, the medium was replaced with NutriStem® containing 3.5 μM CHIR and 500 nM RA (Millipore Sigma, R2625) for RUES2-GLR, and NDiff227 medium containing 10 ng/ml FGF2 for iPS11-mScarlet. On day 4, RUES2-GLR and iPS11-mScarlet cells were subjected to induction.

#### 2. Aggregation of anterior and posterior spheres separately

For the AP gastruloid induction, pre-treated cells were dissociated by StemPro Accutase at 37 °C for 4 mins, counted, and seeded into U-bottom 96-well plates (Nunclon™ Sphera™ 96-Well, Nunclon Sphera-Treated, U-Bottom Microplate, Thermo, 174929) at a density of 2,000 cells per well in 25 μl Essential 6 medium (Thermo, A1516401) containing ROCK inhibitor CEPT (1:200), with 1 μM CHIR and without CHIR, respectively. CEPT was made by mixing 50 nM Chroman 1 (Tocris, 7163), 5 µM Emricasan (Selleck Chemicals, S7775), Polyamine supplement (Millipore Sigma, P8483-5ML) diluted 1:1000, 0.7 µM Trans-ISRIB (Tocris, 5284). After the addition of cells, U-bottom 96-well plates were briefly spun down at 100xg for 1 min and incubated in the 37 °C, 5% CO_2_ incubator for 3 hours.

#### 3. Assembly of anterior and posterior spheres and AP gastruloids culture

At 3 hours, both posterior and anterior cells formed an aggregate. Posterior aggregate was carefully transferred to wells that contain an anterior aggregate to make a total of 50 μl of Essential 6 medium. At 24 hours, an additional 150 μl Essential 6 medium was added to each well. At 48 hours, 150 μl of the medium was carefully removed from each well without disturbing the gastruloid and was replaced with fresh 150 μl of Essential 6 medium supplemented with 100 nM RA and 5% Matrigel. All gastruloids were maintained in 37 °C, 5% CO_2_ incubator throughout the induction up to 120 hours.

### Screening of AP gastruloids induction conditions

For the exploration of the optimum AP gastruloids induction conditions (**Extended Data Fig.1**), 20,000 cells of iPS11-mScarlet and iPS11-eGFP lines were seeded on Vitronectin-coated 12-well plate per well in NutriStem® medium with 10 µM Y-27632 day 0 respectively. Medium for iPS11-mScarlet was replaced by NDiff227 medium (Takara, Y40002) with 10 ng/ml FGF2, 2 μM XAV 939 (STEMCELL technologies, 72672), 0.5 μM LDN193189 (STEMCELL technologies, 72149), and 10 μM SB431542 (Selleck Chemicals, S1067) as indicated on day 2 and day 3. The iPS11-eGFP line was subjected to the posterior induction by NutriStem® medium containing 4 μM CHIR on day 2 and 4 μM CHIR + 500 nM RA on day 3. For induction, 2,000 cells from each line were seeded into each well of U-bottom 96-well plates in 25 μl Essential 6 medium containing 1 μM CHIR. The two aggregates were combined at 0, 3, and 24 hours as indicated. At 24 hours, 150 μl of Essential 6 medium without any supplements was added to each well. At 48 hrs, 150 μl of medium was carefully removed from each well without disturbing the gastruloid and replaced by 150 μl of Essential 6 medium supplemented with 100 nM RA and 5% Matrigel. All gastruloids were maintained in a 37 °C, 5% CO_2_ incubator throughout the induction for 120 hours, and images were taken at the indicated time points.

### Perturbation of human AP gastruloids

For the perturbation of folic acid and ROCK, human AP gastruloids were treated with 10 µM Aminopterin (Millipore Sigma A5159-10VL), 10 µM Y-27632 from 24 hrs to 120 hrs. AP gastruloids were harvested at 72 and 120 hrs for immunostaining and analyses.

### Immunostaining of human AP gastruloids

Gastruloids were collected by wide-bore tips from a 96-well plate at the indicated time points, washed with DPBS twice, and fixed in 4% paraformaldehyde (Electron Microscopy Sciences, 157-4-100) in 12-well plates overnight at 4 °C. The next day, paraformaldehyde was removed and gastruloids were washed by blocking buffer (DPBS containing 0.1% BSA [Gibco, 15260-037], 0.3% Triton X-100 [Sigma, T8787-50ML]) three times and incubated in blocking buffer overnight at 4 °C. Gastruloids can be stored in the blocking buffer at 4 °C for several weeks.

For whole-mount immunostating, fixed gastruloids were incubated in primary antibodies (diluted in blocking buffer at 1:100) overnight at 4 °C, washed in blocking buffer for 3 times and 1 hour each time, then incubated in blocking buffer containing secondary antibodies (1:500-1:1,000 dilution) and DAPI (1:1,000 dilution) overnight at 4 °C and washed in blocking buffer for 3 times and 1 hour each time at dark. For Index matching, gastruloids were transferred to 96-well plates and incubated in 100 μl of CUBIC-R buffer (TCI, T3741) for at least 10-15 mins at room temperature. All samples were analyzed with the confocal microscope (Nikon, Eclipse Ti2). All antibodies used in this study are listed in **Supplementary Table 4**.

All images were analyzed with Fiji^92^ and imageJ^93^. Maximum projections were performed by the function of Z project. The cross-section views were obtained by the function of Orthogonal views, and the position was indicated as a dashed line or numbered arrows in each image.

### Live cell and stereo imaging

For live cell imaging, gastruloids were induced in U-bottom 96-well plates in the 37 °C, 5% CO_2_ incubator with CellCyteX and imaged with an interval of 2 hours for three channels (Brightfield, GFP, and RFP). All images were analyzed with Fiji and imageJ. The length of gastruloids was defined as the largest distance throughout the AP axis, shown in **Extended Data Fig. 2b**. The number of somites was defined by the clear segmentations between somites as indicated by red arrows in **Extended Data Fig. 2d**. The size of somites was measured in the immunostained AP-gastruloid images. Somites were stained with anti-MEOX1 antibody. The diameter was measured by the longest axial of the MEOX1+ cyst across all stacks.

For stereoscope imaging (**Fig. 1c**), gastruloids from three conditions (A-only, P-only, and AP gastruloid) were harvested at 120 hours and fixed in paraformaldehyde in 12-well plates as mentioned in the section of Immunostaining, and then imaged with the stereo microscope.

### Reverse Transcription quantitative Polymerase Chain Reaction (RT-qPCR) of AP gastruloids

For RT-qPCR, 8 AP gastruloids were harvested at 120 hrs, washed twice in DPBS, and manually dissected into three pieces (Anterior, Middle, and Posterior) by microrazer under the microscope. Each region was pooled individually, and its RNA was extracted with PicoPure™ RNA Isolation Kit (Thermo Fisher, KIT0204) according to the manufacturer’s instructions. 500 ng of total RNAs from each sample were used for reverse transcription by High-Capacity cDNA Reverse Transcription Kit (Thermo Fisher, 4374967). The qPCR reaction was run on QuantStudio 6 Flex (Applied Biosystems) by 4 μl of cDNAs (diluted in water by 1:40) mixed with 5 μl of SYBR mix and 1 μl of forward and reverse primers of targeted genes (final concentration 500 nM). The GAPDH was used as the housekeeping gene control. Sequences of primers were obtained from the previous study of neural models^32^ and listed in **Supplemental Table 5**. For analysis, the expression of each gene was first normalized to that of GAPDH in each sample, and then further normalized across samples (i.e., anterior, middle, and posterior) by subtracting the mean and dividing by the standard deviation.

### Single-cell RNA-seq of AP gastruloids by the 10x genomics platform

For cell collection, human AP gastruloids (n=8) were harvested from the 96-well plate at 120 hours and pooled into a 15 ml centrifuge tube, washed with DPBS twice, and dissociated with 1ml 0.05% trypsin-EDTA (Thermo Fisher, 25300054) for 8 mins at 37 °C and quenched with 200 μL FBS. Gastruloids were dissociated into single cells by pipetting, pelleted by centrifuging at 300 rcf for 3 mins, resuspended in StemCell Banker (Amsbio, 11924), and stored at −80 °C.

Single-cell RNA-seq (scRNA-seq) libraries were created using 10x Genomics Chromium platform, GEM-X Universal 3’ Gene Expression v4 reagents kit following manufacturer specifications and guidelines. In brief, dissociated and frozen single cell suspension were thawed in Essential 6 medium containing 10% FBS, washed with PBS with 0.04% BSA, counted and diluted to 1500 cell/uL to target an optimal recovery of 20,000 cells/lane. The generated scRNA-seq libraries were visualized and quantified by TapeStation (Agilent), and sequenced on NextSeq 2000 (Illumina) with the following read structure: Read 1: 28 cycles; i7 Index: 10 cycles; i5 Index: 10 cycles; Read 2: 90 cycles.

### Single-cell RNA-seq data processing for the AP gastruloids

The single-cell transcriptome data from AP gastruloids were processed using Cell Ranger 8.0.0^94^ with default settings and the reference genome refdata-gex-GRCh38-2020-A. From the “raw_feature_bc_matrix” folder, we extracted the gene-by-cell matrix, filtering out cells with fewer than 1,000 UMIs or 500 detected genes. Genes from chromosomes 1-22, X, Y, and MT were retained. Doublets were identified using Scrublet/v0.1^95^, and we calculated the percentage of reads mapping to mitochondrial (MT%) and ribosomal (Ribo%) genes for each cell. After manually examining the distribution of UMIs, MT%, and Ribo% across cells, we applied criteria to further remove potentially low-quality cells: those with UMI counts fewer than 5,000 or exceeding the top 0.1% of the total cells, doublet scores calculated by Scrublet over 0.2, MT% over 10%, or Ribo% over 40%, were removed. A total of 13,058 cells were retained.

Next, we applied Seurat/v5^61^ to perform an initial round of dimensionality reduction and cell clustering: 1) retaining protein-coding genes and lincRNA for each cell and removing gene counts mapping to sex chromosomes; 2) normalizing UMI counts by the total counts per cell, multiplying by a pseudocount of 10,000, and applying log transformation; 3) selecting the 2,500 most highly variable genes, scaling their expression to a zero mean and unit variance, and regressing out the cell cycle index calculated using the CellCycleScoring function; 4) applying PCA (dims = 30); 5) using the top 30 corrected PCs to calculate a neighborhood graph, followed by louvain clustering (resolution = 1); 6) performing UMAP visualization in 2D space (min.dist = 0.3). Of note, we identified two cell clusters enriched for cells with substantially higher doublet scores compared to others. We removed cells from these clusters (∼4.5% of the total) and repeated steps 1–6, resulting in a final dataset of 12,469 cells. By manually reviewing canonical cell-type-specific marker genes (**Supplementary Table 1**), we merged adjacent clusters as needed, annotating 11 major cell types. To further refine cell type annotations, we performed subclustering on four subsets of cells: I) MHB and hindbrain, II) neural tube and early neurons, III) neural crest and head mesoderm, and IV) somites and intermediate mesoderm. Incorporating these subcluster annotations with the major cell types, we ultimately identified a total of 27 distinct cell types.

### Integration of AP gastruloids with other datasets

To validate cell type identification and computationally infer the developmental stage of AP gastruloids, we integrated them with several publicly available datasets. First, we compared them with natural mouse embryos, including two published comprehensive datasets profiling gastrulation^45^ (E6.5–E8.5; n = 108,857 cells) and organogenesis^7^ (E8–P0). For the organogenesis dataset, cells from E8–E10 (early somitogenesis) were selected and randomly downsampled to 100,000. Second, we integrated with recently published RA gastruloids^4^ (n = 18,324 cells, 120 h). Third, we included primate datasets, comprising CS8–CS11 monkey embryos^59^ (n = 56,636 cells) and CS12–CS16 human embryos^16^ (n = 185,140 cells). Integration across two or more datasets was performed using Seurat’s integration workflow^61^: FindIntegrationAnchors was applied to identify shared anchors, followed by IntegrateData for batch correction and alignment. Cross-species integration was facilitated by mapping homologous genes using BioMart, retaining many-to-many pairs.

To infer the developmental stage of AP gastruloids *in silico*, based on the integrated dataset of AP gastruloids with monkey and human embryo datasets, for each cell from the human AP gastruloid or from each embryonic stage (CS8, CS9, and CS11 for monkey; CS12, CS13–14, and CS15–16 for human), the 15 nearest neighbors were identified in the 30-dimensional PC space within each corresponding embryonic stage. The distribution of these *in vivo* nearest neighbors was then collectively used to infer the potential developmental stage of AP gastruloids or of individual AP gastruloid cell types.

### Identification of correlated rhombomere between AP gastruloids and mouse

To identify correlated cell types of the MHB and hindbrain (r1–r8) between AP gastruloids and mouse embryo organogenesis datasets (E8–E8.5 only), we first normalized gene expression in each cell by the total expression, multiplied by a pseudocount of 10,000, and applied log transformation. We then calculated the average expression of each gene across all cells within each cell type. To account for differences in cell numbers across cell types, we randomly downsampled AP gastruloid cells to 200 per cell type and mouse embryo cells to 500 per cell type. Next, for the two datasets, we applied non-negative least squares (*NNLS*) regression to predict gene expression in a target cell cluster (*T_a_*) in dataset A based on the gene expression of all cell clusters (*M_b_*) in dataset B: *T_a_ = β_0a_ + β_1a_M_b_*, based on the union of the 1,500 most highly expressed genes and 1,500 most highly specific genes in the target cell cluster. We then switched the roles of datasets A and B, *i.e.,* predicting the gene expression of target cell cluster (*T_b_*) in dataset B from the gene expression of all cell clusters (*M_a_*) in dataset A: *T_b_ = β_0b_ + β_1b_M_a_*. Finally, for each cell cluster *a* in dataset A and each cell cluster *b* in dataset B, we combined the two correlation coefficients: *β* = 2(*β_ab_* + 0.01)(*β_ba_*+ 0.01) to obtain a statistic, where high values reflect reciprocal, specific predictivity.

### Assignment of AP gastruloid cells to hPSC origins

Cells of AP gastruloids were assigned to their origins from two hPSC lines, iPSC11-mScarlet (anteriorized) and RUES2-GLR (posteriorized), using Vireo^60^ with default settings without the reference SNP genotypes (https://demultiplexing-doublet-detecting-docs.readthedocs.io/en/latest/Vireo.html#vireo-docs). Vireo is a Bayesian method that demultiplexes pooled scRNA-seq data by inferring donor genotypes from allelic counts, allowing cell assignment and doublet detection without external genotype references. Based on this analysis, 61% of cells were assigned to the anteriorized iPSC11-mScarlet line, 36% to the posteriorized RUES2-GLR line, 2% were identified as doublets, and 1% remained unassignable.

---

**Extended Data Fig. 1.**
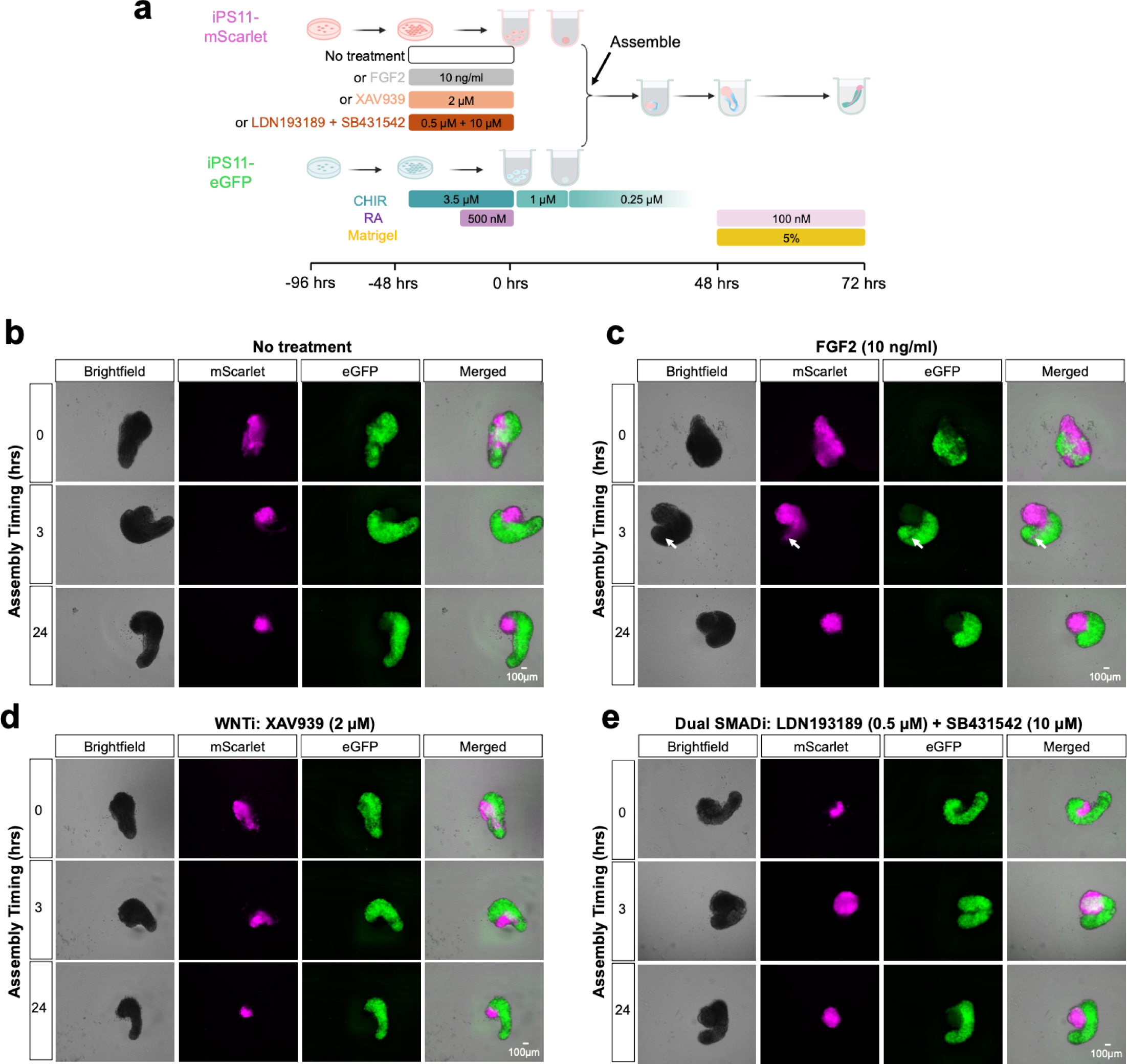
Assembly dynamics of anteriorized and posteriorized hPSCs. **(a)** We sought to evaluate the impact of varying the anteriorizing signal and the timing of assembly. For these experiments, we used hPSC lines tagged with mScarlet (anteriorized) or eGFP (posteriorized) respectively. iPS11-mScarlet cells were subjected to pre-treatment with one of three possible anteriorizing cocktails for 2 days (−48 hrs to 0 hrs): FGF2, XAV939 (a WNT inhibitor) or a combination of LDN193189 and SB431542 (dual SMAD inhibition). iPS11-eGFP cells were subjected to pre-treatment with a posteriorizing cocktail (CHIR→CHIR+RA)^4^ for 2 days (−48 hrs to 0 hrs). These two populations were either mixed as disaggregated cells at 0 hrs or assembled as aggregates at 3 hrs or 24 hrs. **(b-e)** Representative images of the resulting gastruloids under various conditions at 72 hrs. Each panel represents a distinct combination of anteriorizing pre-treatment and assembly timing. Scale bar: 100 μm. **(b)**, no anteriorizing pre-treatment; **(c)**, 10 ng/ml FGF2; **(d)**, 2 µM XAV939; and **(e)**, 0.5 µM LDN193189 and 10 µM SB431542. Rows correspond to different assembly timings as indicated on the left (0, 3 or 24 hrs), with the same field of view shown four times (from left to right: brightfield, mScarlet, eGFP, merged). In panel **c**, the identically positioned white arrows in the row of images corresponding to 3-hr assembly highlight the intercalation of anteriorized mScarlet+ cells into posteriorized eGFP+ cells.

**Extended Data Fig. 2.**
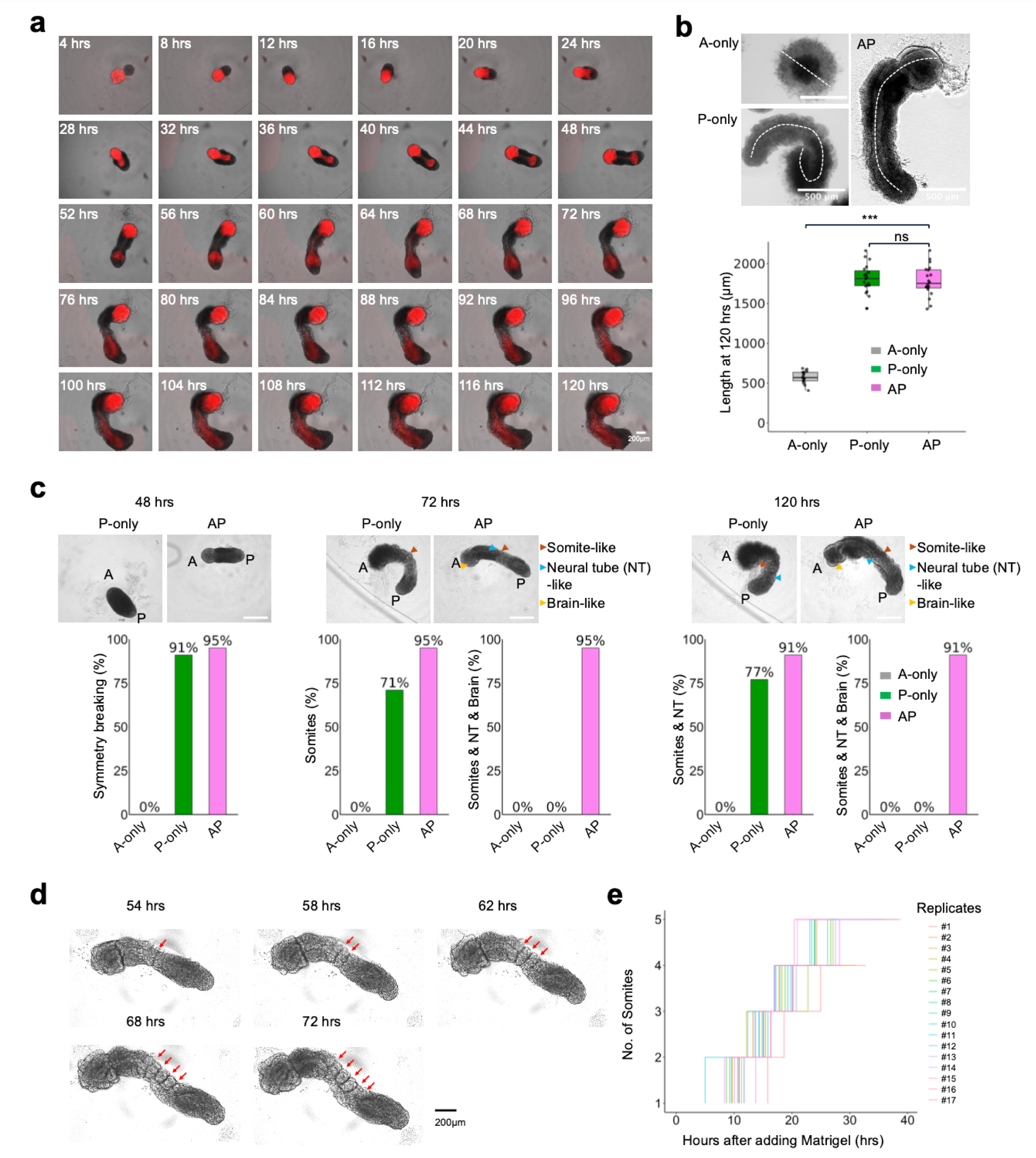
Live cell imaging and morphological characterization of human AP gastruloids. **(a)** Live cell imaging of a representative AP gastruloid from 4 to 120 hrs, with 4-hr intervals. All images shown merge brightfield and mScarlet (red) channel views to highlight how anteriorized mScarlet+ iPS11 cells ingress towards posteriorized RUES2-GLR cells (mScarlet-) to contribute to a seemingly continuous neural tube-like structure along the AP axis. Scale bar: 200 μm. **(b)** Length distributions of A-only aggregates, P-only gastruloids (*i.e.* RA gastruloids^4^) and AP gastruloids at 120 hrs. Left: Boxplot of lengths of A-only aggregates (n=15), P-only gastruloids (n=19) and AP gastruloids (n=20). Thick horizontal lines, median; upper and lower box edges, third and first quartiles, respectively; whiskers, largest to smallest values within 1.5 times the interquartile range. Dots beyond the whiskers: outliers. Student’s t-test was used for statistical tests. ns: not significant. ***: p_value<0.001. Right: Illustration of how lengths were assessed. For A-only aggregates, diameters were used as no elongation was observed. For P-only and AP gastruloid, the longest distance along the midline along the AP axis was used as the length. Scale bar: 500 μm. Student’s t-test was used for statistical tests. ns: not significant. ***: p-value<0.001. **(c)** Morphological characterization at 48 hrs (left), 72 hrs (middle) and 120 hrs (right). Top row: representative images of P-only and AP gastruloid, with arrowheads marking our annotation of specific structures. AP axis is indicated by A and P labels. Brown, blue and yellow arrowheads annotate somite, neural tube and brain-like structures, respectively. Bottom row: Barplots of percentages of examined aggregates/gastruloids exhibiting specific structures at specific timepoints. No elongation, symmetry breaking or advanced embryo-like structures were observed in any A-only aggregates (0%; n=35 for all timepoints). The vast majority of both P-only (91%, n=46) and AP (95%, n=87) gastruloids exhibited modest elongation and differential cellular morphologies at their A and P poles (symmetry breaking) by 48 hrs. Segmented somites were observed in 71% of P-only gastruloids (n=49) by 72 hrs. In 77% of P-only gastruloids (n=52), both somites and a neural tube-like structure were evident by 120 hrs. In sharp contrast with A-only aggregates and P-only gastruloids, nearly all AP gastruloids contained all three structures (somite, neural tube, and brain-like) at 72 hrs (95%; n=77), and these structures were overwhelmingly maintained through to the 120 hr timepoint (90.9%, n=77). **(d-e)** Quantification of segmentation clock in AP gastruloids. In this experiment, Matrigel was added at 44 hrs and AP gastruloids (n=17) were subsequently imaged every 2 hrs. **(d)** Representative images of a single AP gastruloids at 54, 58, 62, 68 and 72 hrs. Red arrows point to the border of segmented somites. **(e)** The timings of segmentation of new somites are shown as the number of hours (hrs) after the addition of Matrigel (*x*-axis) and the total number of somites is shown along the *y*-axis. Each line corresponds to an individual AP gastruloid from #1-#17.

**Extended Data Figure 3.**
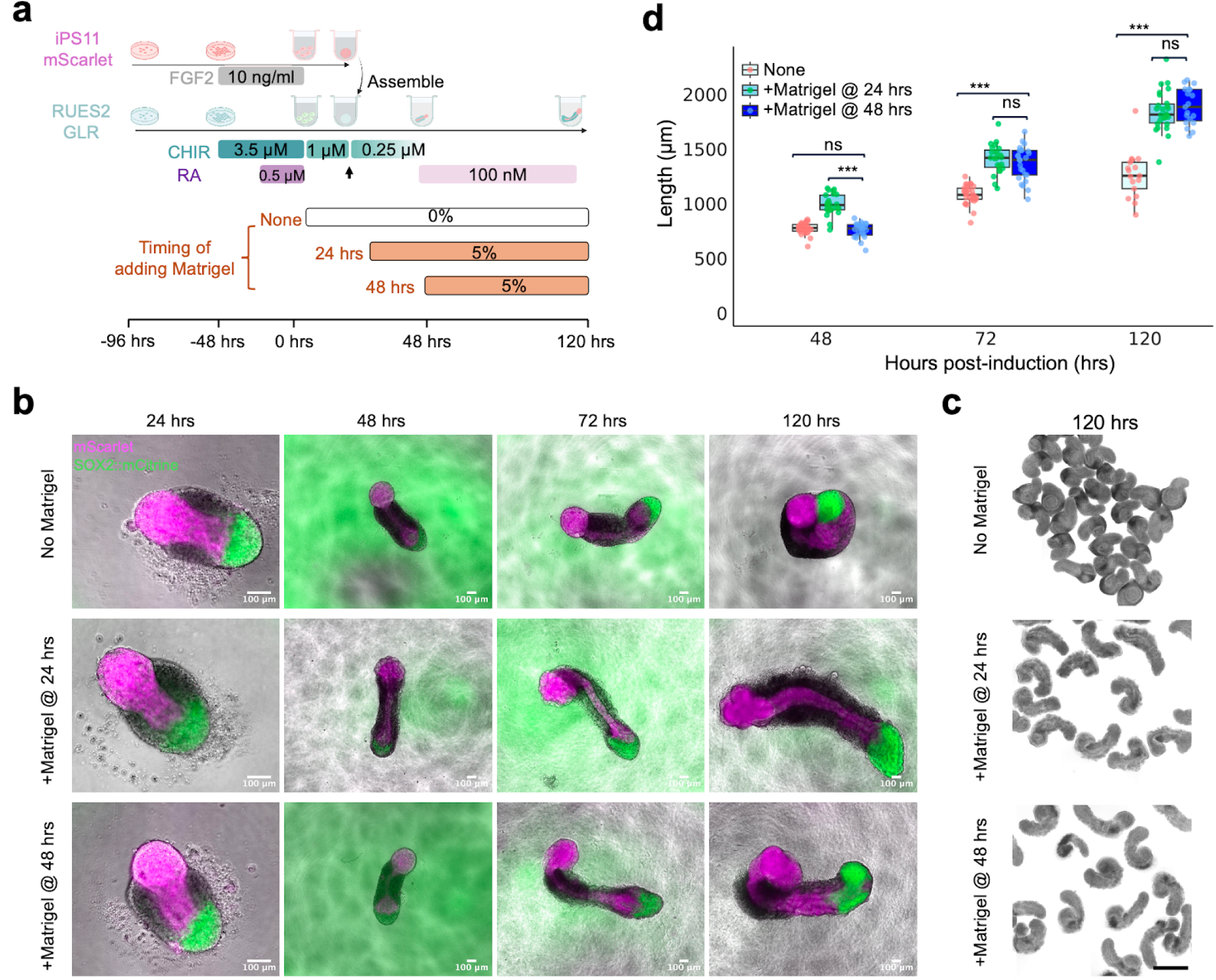
Impact of withholding or accelerating Matrigel supplementation on AP gastruloid morphology. **(a)** Schematic of experiment in which three conditions were tested: no Matrigel supplementation at any timepoint (*i.e.* withholding), adding Matrigel at 24 hrs (*i.e.* accelerating) or adding Matrigel at 48 hrs (*i.e.* baseline protocol). **(b)** Representative images of the resulting gastruloids from 24 to 120 hrs under the three conditions tested. In these merged images, magenta corresponds to mScarlet signal from anteriorized iPS11-mScarlet cell line, and green to SOX2::mCitrine signal from the posteriorized RUES2-GLR cell line. Scale bar: 100 μm. **(c)** Brightfield images of gastruloids collected at 120 hrs under the three conditions tested. Scale bar: 500 μm. **(d)** Boxplots of gastruloid lengths, under the three conditions tested, measured at 48, 72 and 120 hrs. Pink, green and blue correspond to ‘no Matrigel’, ‘adding Matrigel at 24 hrs’, and ‘adding Matrigel at 48 hrs’ conditions, respectively. The number of gastruloids measured per condition per timepoint ranged from 17-29. Thick horizontal lines, median; upper and lower box edges, third and first quartiles, respectively; whiskers, largest to smallest values within 1.5 times the interquartile range. Dots beyond the whiskers: outliers. Student’s t-test was used for statistical tests. ns: not significant. ***: p_value<0.001.

**Extended Data Figure 4.**
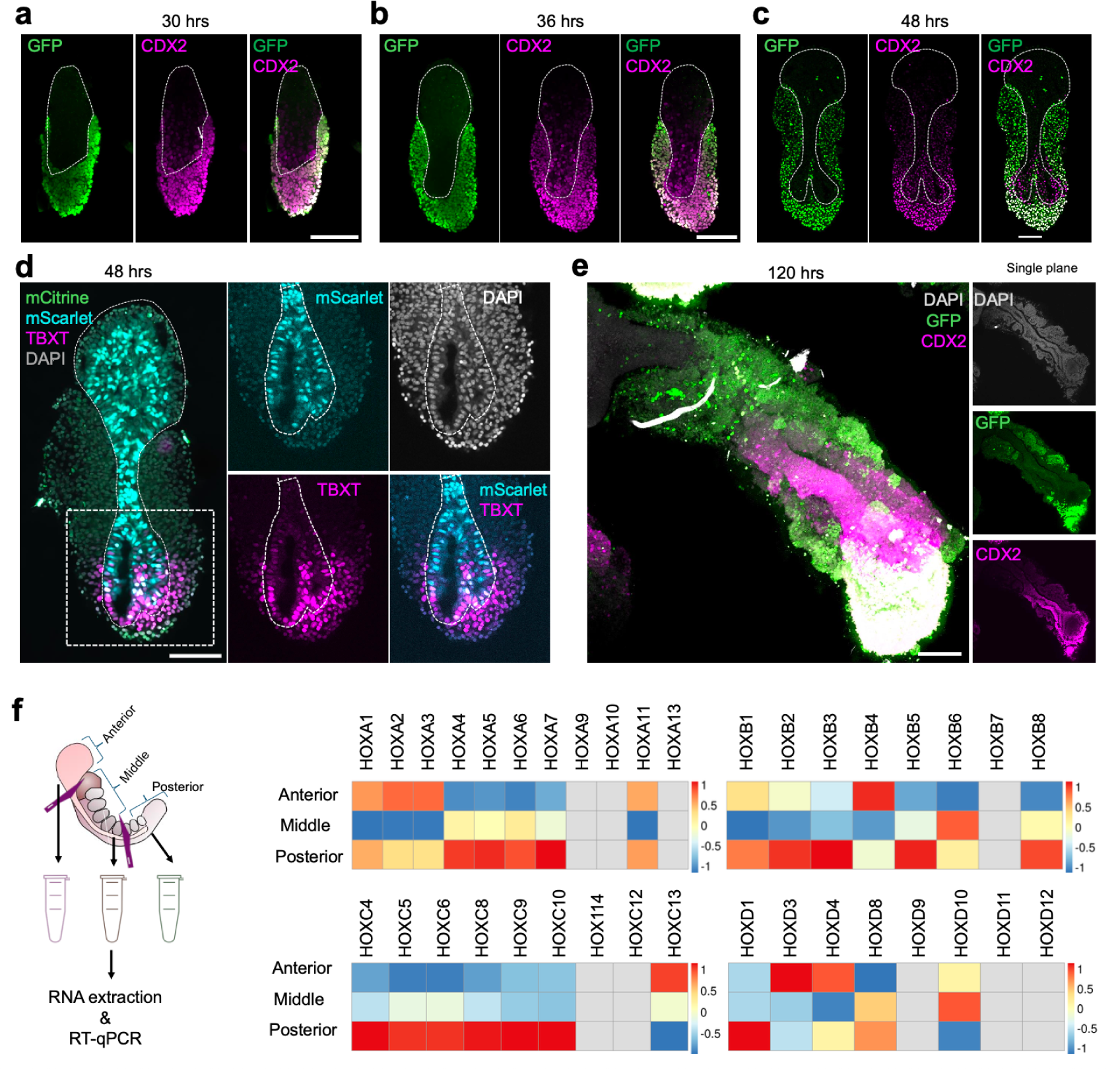
AP axis patterning in human AP gastruloids. **(a-c)** Time-course immunostaining of AP gastruloids with GFP (green) and CDX2 (magenta) antibodies at 30, 36 and 48 hrs. All images are displayed as single planes and white dashed outlines highlight regions derived from the anteriorized cells. The GFP antibody recognizes both SOX2::mCitrine and TBXT::mCerulean. CDX2+/GFP-cells are inferred to correspond to NMPs derived from anteriorized iPS11-mScarlet hPSCs. Scale bar: 100 μm. **(d)** Images from one AP gastruloid at 48 hrs stained with DAPI (gray) and TBXT (magenta) antibody. mScarlet (cyan) and mCitrine (green) indicate retained mScarlet and SOX2::mCitrine proteins in anteriorized iPS11-mScarlet derived cells and posteriorized RUES2-GLR derived cells, respectively. Left: a single plane with all channels merged. Middle and right: Enlarged single channel views of a subregion (white dashed rectangle). White dashed outlines highlight regions derived from the anteriorized cells. TBXT+ cells within the white dashed outline are inferred to correspond to NMPs derived from anteriorized hPSCs, while TBXT+ cells outside the white dashed outline are inferred to correspond to NMPs derived from posteriorized hPSCs. Scale bar: 100 μm. **(e)** Images from one AP gastruloid at 120 hrs stained with GFP (green) and CDX2 (magenta) antibodies and displayed as maximum projection (left) or single plane (right) image. The GFP antibody recognizes both SOX2::mCitrine and TBXT::mCerulean. Single plane DAPI image (top right) showcases the neural tube structure flanked by somites. Notably, caudally located CDX2+ cells are GFP+ while CDX2+ cells located in the posterior neural tube are GFP-. Scale bar: 100 μm. **(f)** Normalized expression of HOX genes along the AP axis obtained by performing RT-qPCR on sectioned AP gastruloids. Left: Experimental schematic. 8 AP gastruloids were sectioned into three pieces (anterior, middle and posterior) at 120 hrs under the microscope and subjected to RNA extraction and RT-qPCR of HOX genes. Right: Heatmaps of the resulting expression pattern of HOX genes. Each column stands for one gene as indicated on the top and each row stands for the sampling location on the left. The expression of each gene was first normalized to that of the housekeeping gene GAPDH in each sample, and then further normalized across samples (*i.e.* anterior, posterior, posterior) by subtracting the mean and dividing by the standard deviation.

**Extended Data Figure 5.**
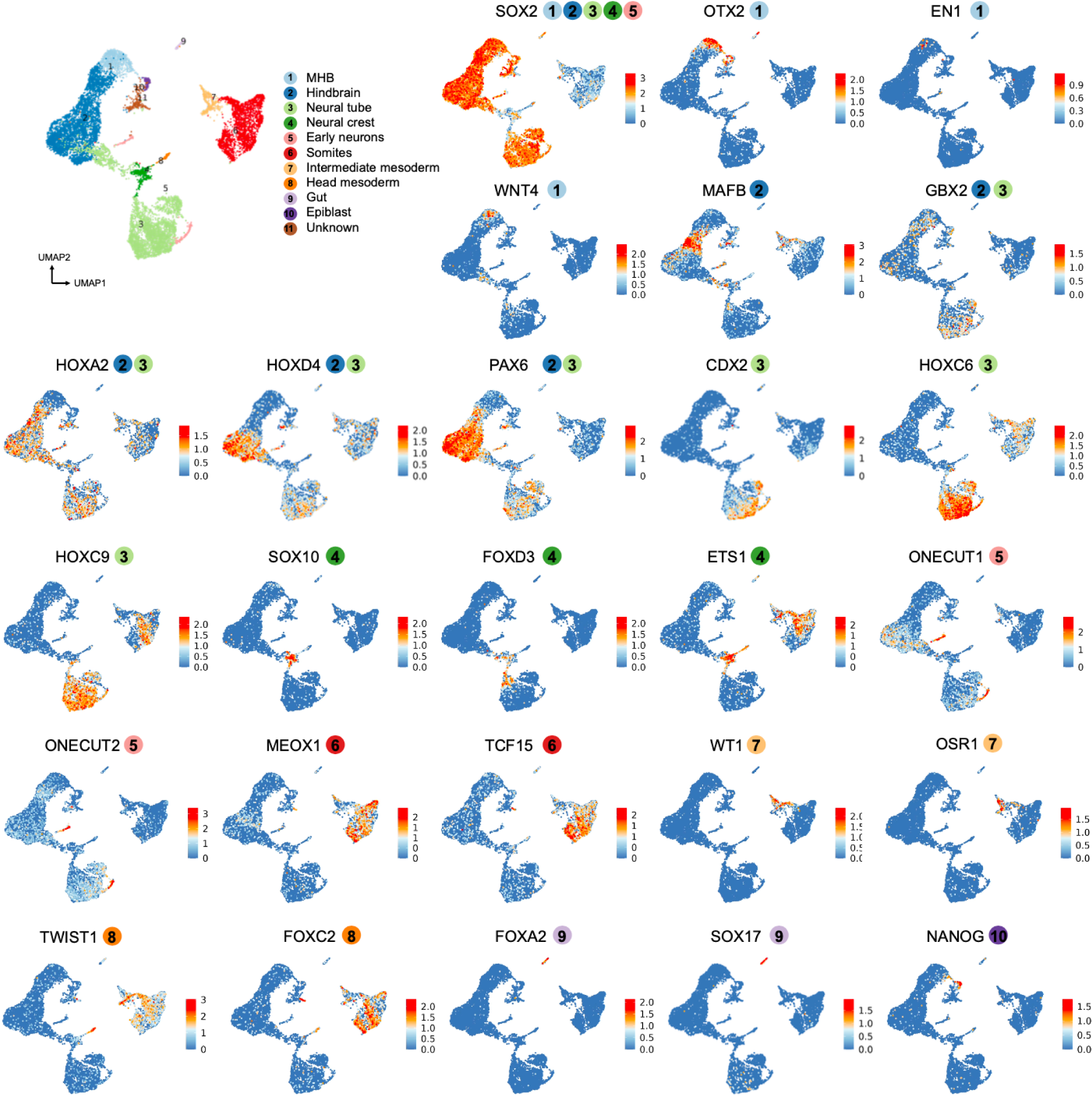
Marker gene-based annotation of main cell types in human AP gastruloids. The same global UMAP as shown in the top left/right panels of Fig. 3a, colored by cell types (top left) or expression of select marker genes (raw UMI counts divided by total counts per cell, multiplied by 10,000, and natural log transformed) that were used to annotate main cell types. The numbered/colored circles next to each gene name indicate which cell type(s) the marker gene corresponds to, with references provided in **Supplementary Table 1**.

**Extended Data Figure 6.**
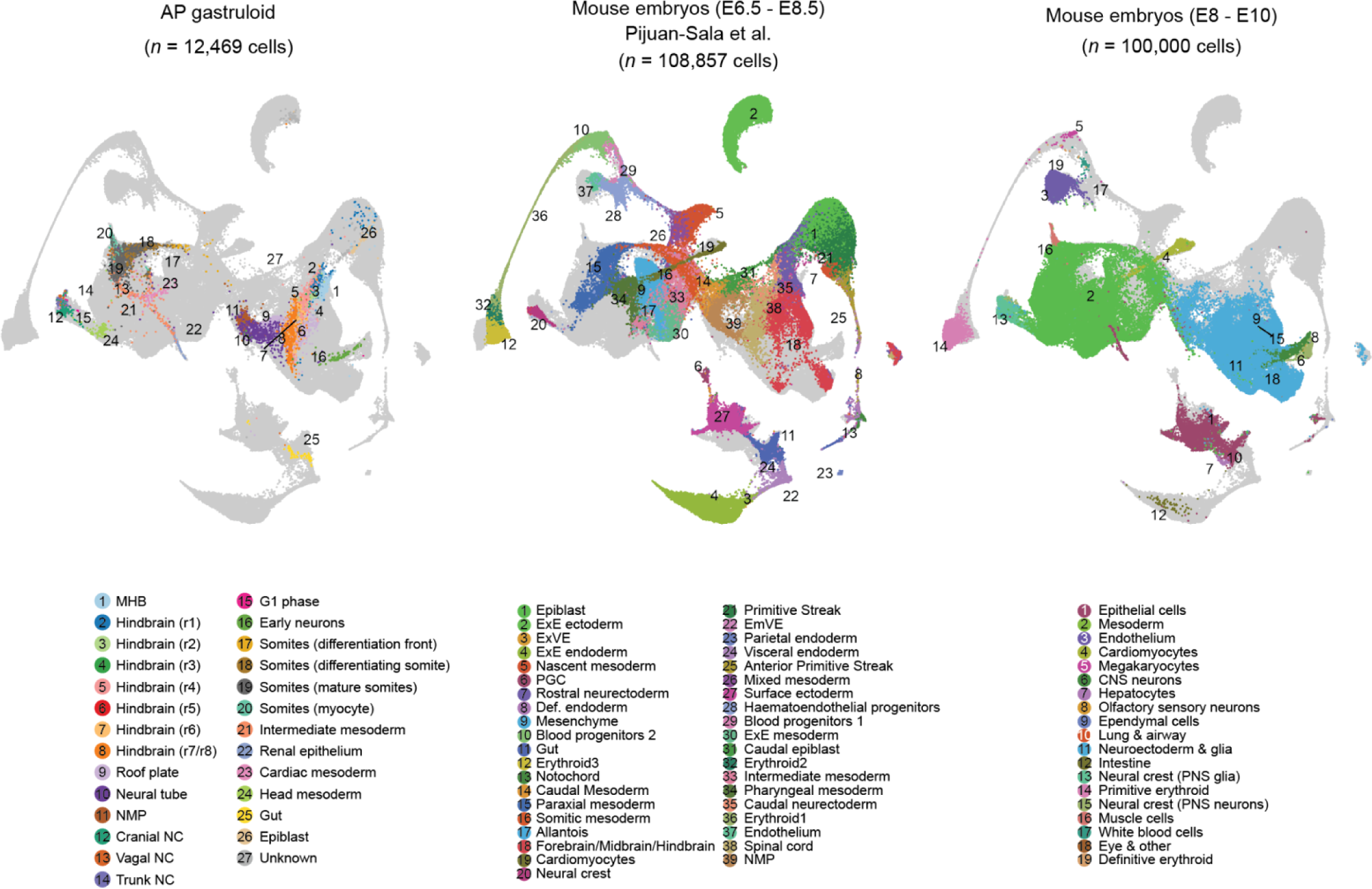
Integrating and co-embedding cells from *in vitro* human AP gastruloids and *in vivo* mouse embryos to validate cell type annotations. UMAP visualization of 221,326 co-embedded cells from three datasets, including AP gastruloids from this study and two studies of mouse embryos during gastrulation (E6.5-E8.5)^45^ or early somitogenesis (E8-E10)^7^, after batch correction^61^. The same UMAP is shown three times, with colors highlighting cells from each dataset. To align with the cell counts in the other datasets, sci-RNA-seq3 profiles from the mouse embryos early somitogenesis^7^ (right) were randomly subsampled to 100,000 cells. To refine cell type annotations, the 20 nearest neighbors of AP gastruloid cells were identified in the PCA space (30 dimensions) from the two embryo datasets. For each AP gastruloid cell type, the proportions of these nearest neighbors across embryo cell types were calculated, and the five most abundant embryo cell types are listed in **Supplementary Table 2**.

**Extended Data Figure 7.**
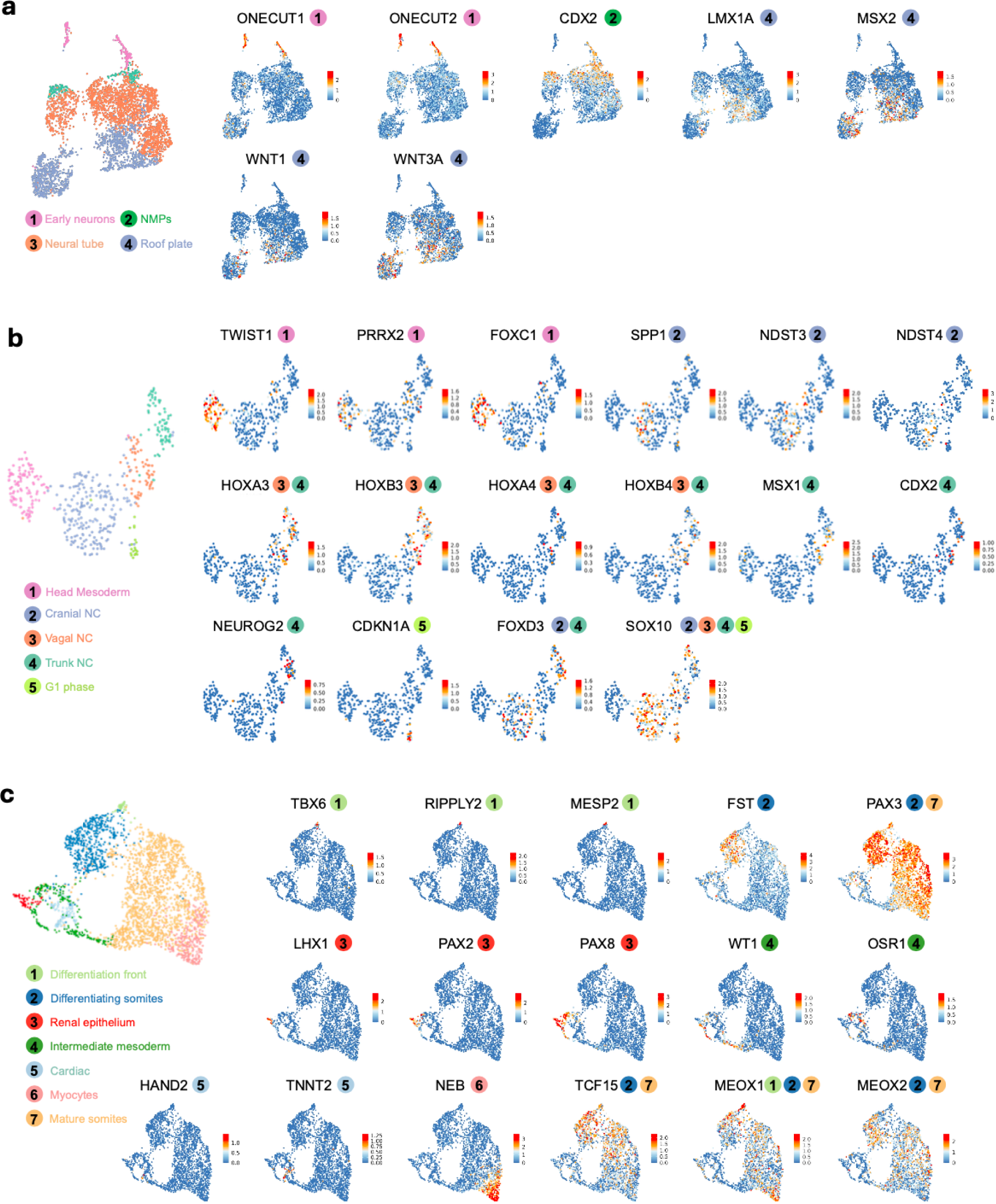
Marker gene-based annotation of cell subtypes in human AP gastruloids. **(a-c)** Identical UMAPs as subpanels II-IV of Fig. 3a, which correspond to **(a)** neural tube/early neurons (II, 3,724 cells), **(b)** neural crest/head mesoderm (III, 464 cells), and **(c)** somitic/intermediate mesoderm (IV, 2,790 cells). These are colored by cell types (left) or expression of select marker genes (raw UMI counts divided by total counts per cell, multiplied by 10,000, and natural log transformed) that were used to annotate cell subtypes. The numbered/colored circles next to each gene name indicate which cell type(s) the marker gene corresponds to, with references provided in **Supplementary Table 1**.

**Extended Data Figure 8.**
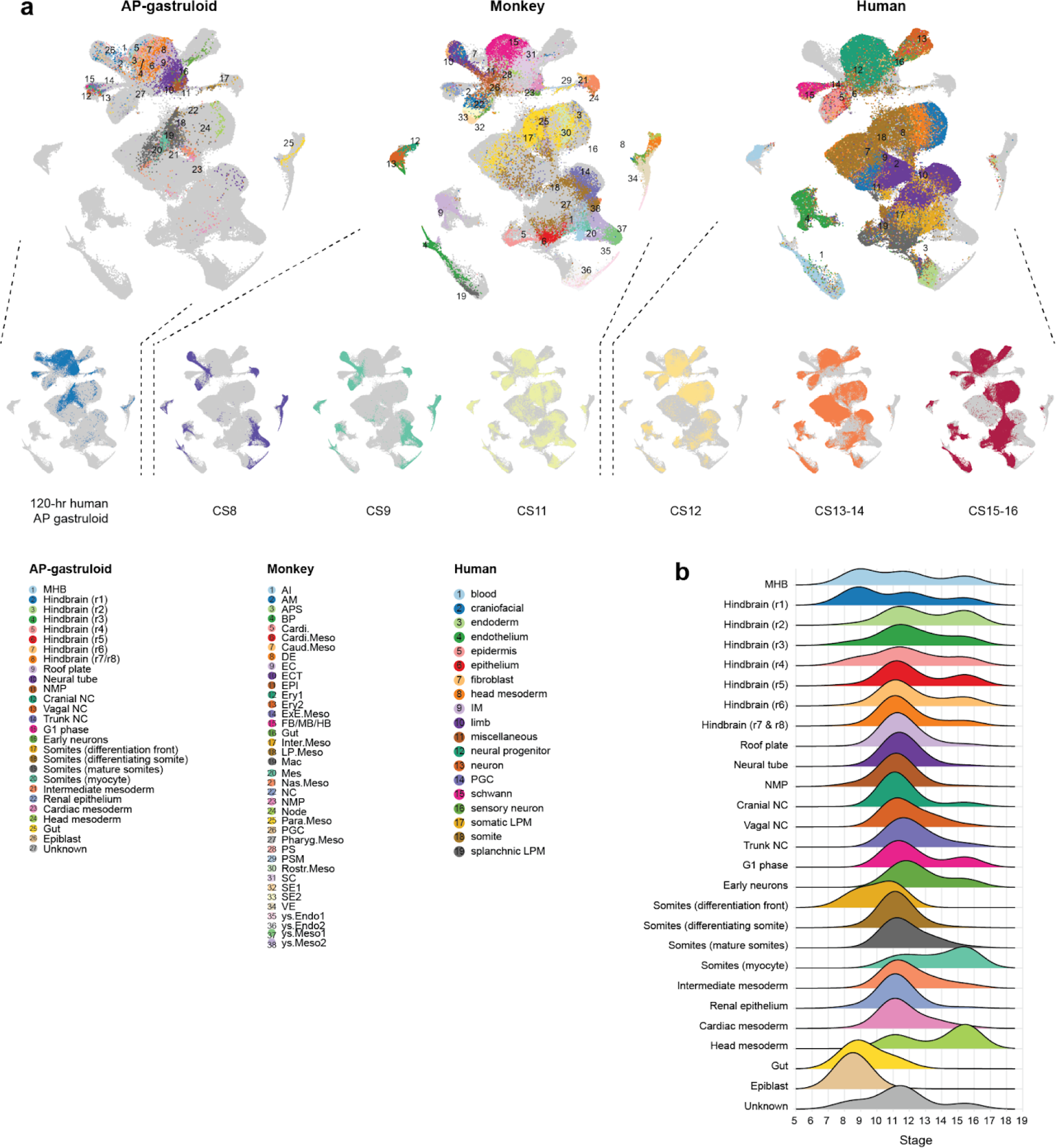
In silico staging of human AP gastruloids. **(a)** UMAP visualization of 254,245 co-embedded scRNA-seq profiles from three datasets: 120-hr human AP gastruloids from this study (12,469 cells), CS8-CS11 monkey embryos (56,636 cells)^59^, and CS12-CS16 human embryos (185,140 cells)^16^, after batch correction^61^. The same UMAP is shown three times to highlight different datasets, with cell types labeled and annotations highlighted on the left. In the right columns, cells from this study and from various stages of monkey and human embryos are highlighted in each UMAP. **(b)** For each cell from the human AP gastruloid or from each human/monkey embryonic stage (CS8, CS9, and CS11 for monkey; CS12, CS13-14, and CS15-16 for human), the 15 nearest neighbors were identified within each human or monkey embryonic stage. The distribution of stages among these 15 *in vivo* nearest neighbors is plotted for each individual cell type. CS13–14 and CS15–16 are represented as 13.5 and 15.5, respectively, on the x-axis. YS endoderm: Yolk sac endoderm. DE: Definitive endoderm. SE: surface endoderm. YS meso: Yolk sac mesoderm. ExMC: Extra-embryonic mesoderm. PSM: presomitic mesoderm. NMP: Neuromesodermal progenitor. NC: neural crest.

**Extended Data Figure 9.**
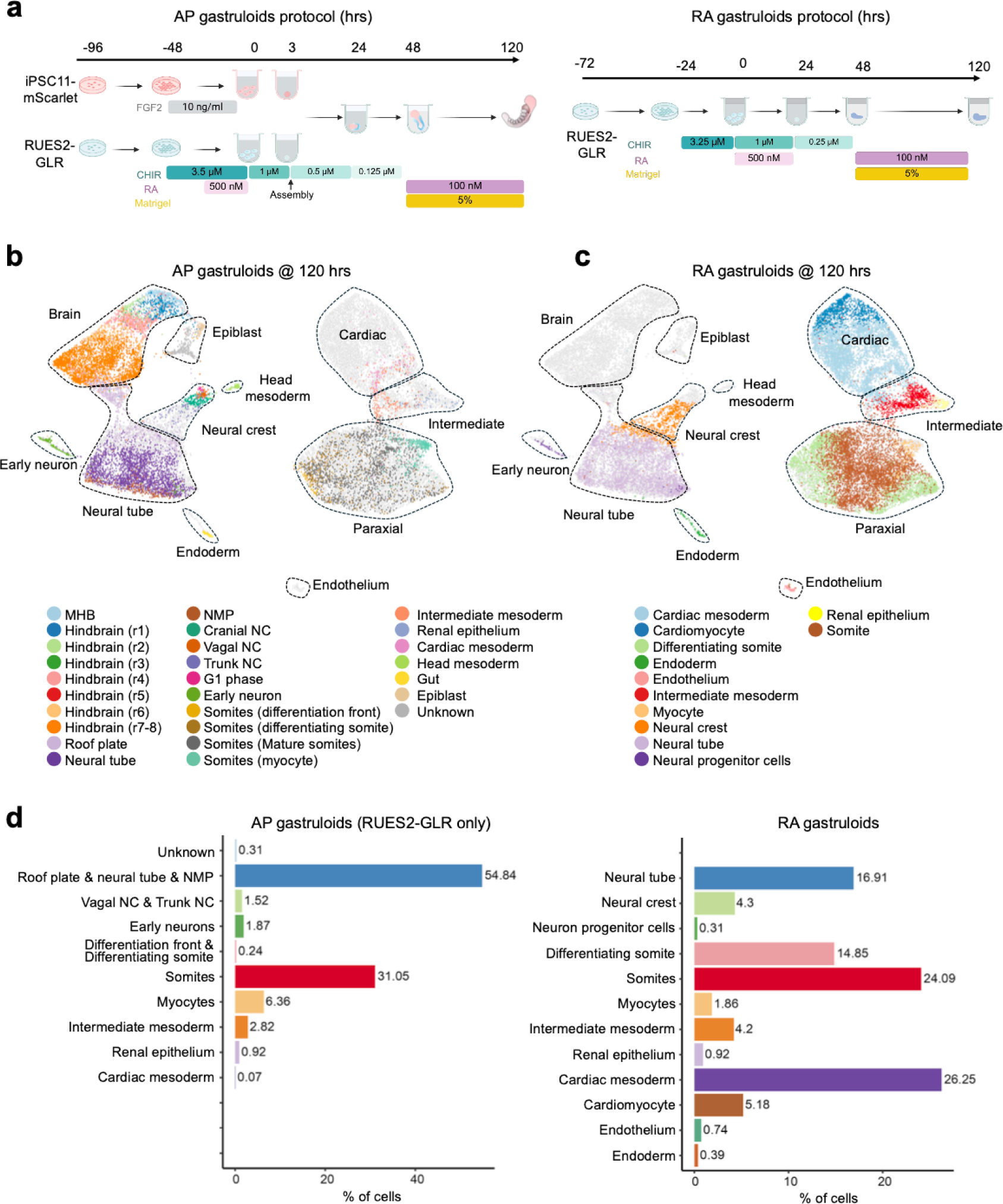
Comparison of human AP gastruloids vs. human RA gastruloids. (**a**) Schematic of protocols for AP (left) and RA gastruloids^4^ (right) induction. In both protocols, RUES2-GLR cells were subjected to the CHIR and RA treatment in the initial induction phase, followed by the additional RA exposure together with the Matrigel supplement during the elongation phase, albeit with some differences in timing and concentration, together with the major difference of the addition of anteriorized iPSC11-mScarlet cells in the AP gastruloid protocol only. (**b-c**) Integration and co-embedding of scRNA-seq datasets from AP gastruloids (this study: 12,469 cells) and RA gastruloids at 120 hrs (GSE208369: 18,324 cells)^4^ with the RPCA method in the Seurat package^61^. All cells were re-clustered in the integrated UMAP but are colored by the original annotation of cell types from AP (panel **b**) or RA (panel **c**) gastruloids. Cells from RA gastruloids are colored by light gray in panel **b** and cells from AP gastruloids are colored light gray in panel **c**. Major cell lineages are manually bounded by dashed polygons and named in panel **b**, except for endothelium which only appears in RA gastruloids and thus is only labeled in panel **c**. (**d**) Barplot of percentages of different cell types derived from the RUES2-GLR line in AP (left) and RA (right) gastruloids. Note that cells inferred to have derived from iPSC-mScarlet hPSCs in AP gastruloids are excluded from this analysis.

**Extended Data Figure 10.**
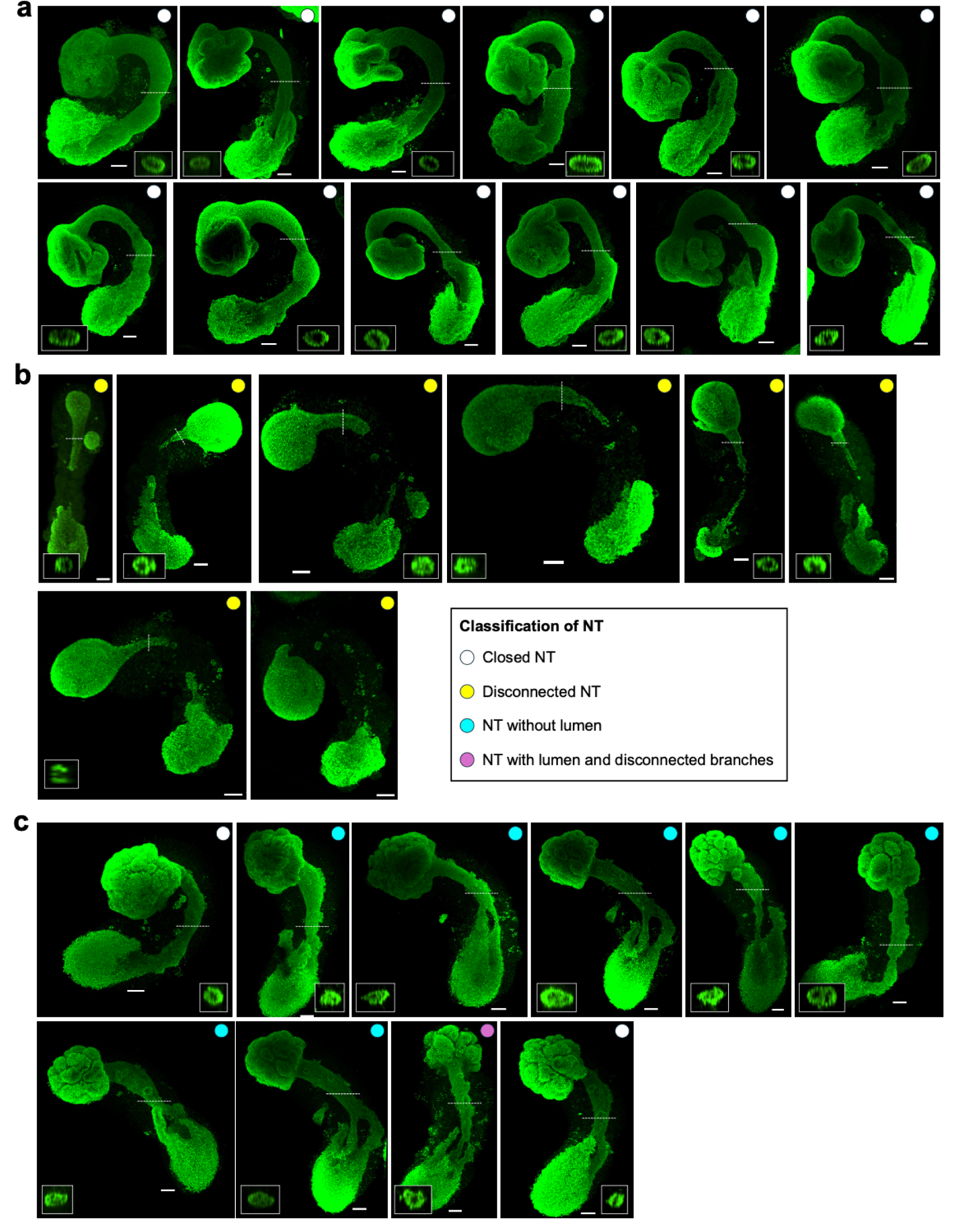
Maximum projections of SOX2 channel (green) of AP gastruloids at 120 hrs under conditions of Ctrl (a), aminopterin (b) and Y-27632 (c). In each image, the cross-sectional image from the position indicated by the dashed white line was embedded in the white rectangle at either bottom corner to highlight the neural tube lumen structure (or lack thereof). Colored dots at the top left corner indicate classifications of neural tube structures. White: successful closed neural tube. Yellow: disconnected neural tube. Cyan: neural tube without lumen. Magenta: neural tube with lumen and disconnected branches.

**Extended Data Figure 11.**
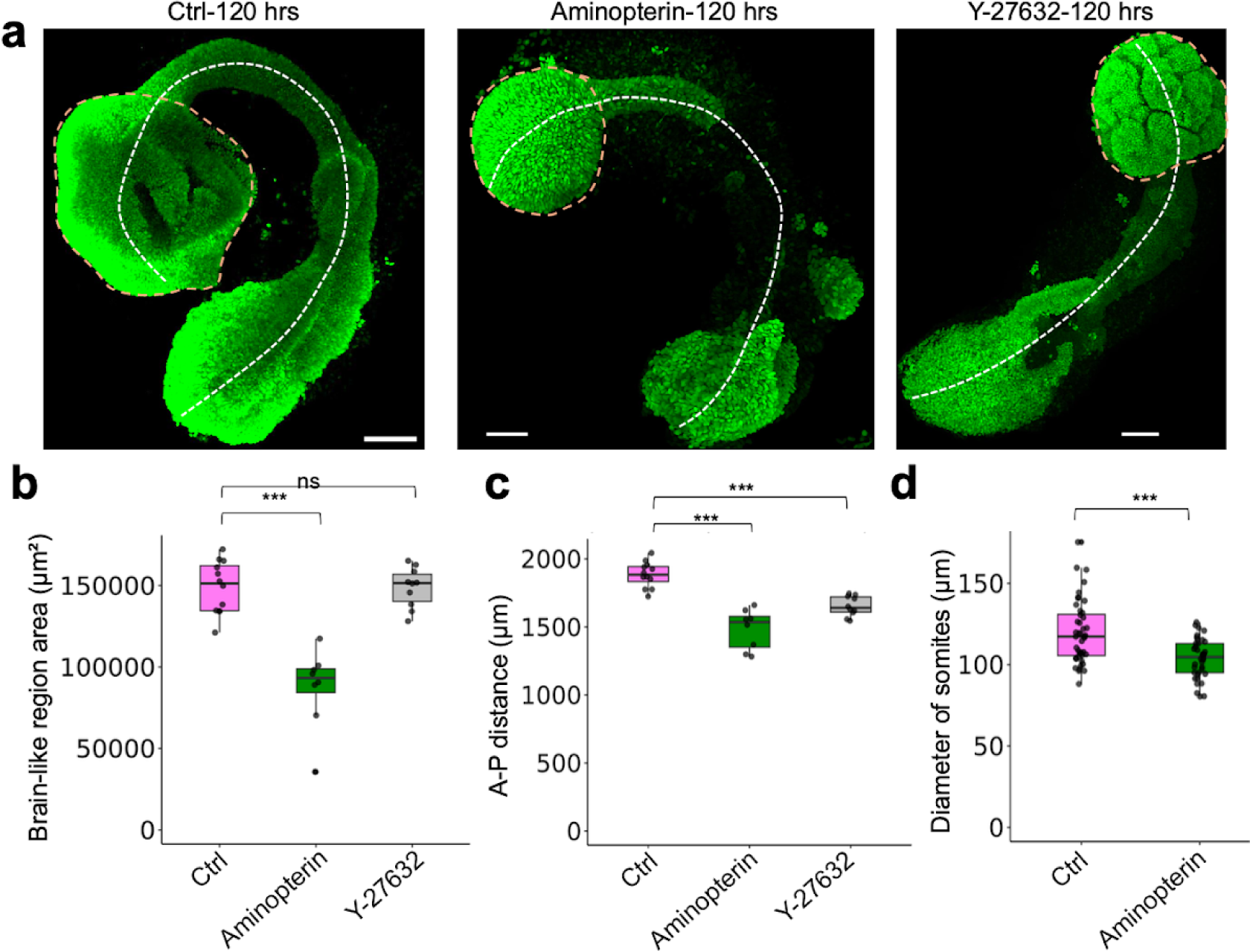
Quantification of morphological features of AP gastruloids under chemical perturbation. (**a**) Maximum projection of same gastruloids at 120 hrs as shown in Fig. 5e**-g**, further highlighting how measurements were performed. Dashed orange circle: measured areas of the brain-like regions. Dashed white line: measured distance of anterior-posterior ends as the overall length of AP gastruloids. (**b**) Box plot of area of brain-like region in each condition. Ctrl: 148,884 ± 16,094 µm², n=12. Aminopterin: 87,237 ± 24,636 µm², n=8. Y-27632: 148,775 ± 12,206 µm², n=10. (**c**) Box plot of the length of AP gastruloids in each condition. Ctrl: 1885 ± 94 μm, n=12. Aminopterine: 1484 ± 147 μm, n=8. Y-27632: 1653 ± 73 μm, n=10. **(d)** Box plot of the diameter of segmented somites in Ctrl and Aminopterin (segmented somites were not observed in Y-27632 condition, as discussed in text). Ctrl: 120 ± 19 μm, n=44 from 5 gastruloids. Aminopterin: 104 ± 12 μm, n=38 from 5 gastruloids. (**b-d**) Thick horizontal lines, median; upper and lower box edges, third and first quartiles, respectively; whiskers, largest to smallest values within 1.5 times the interquartile range. Dots beyond the whiskers: outliers. Student’s t-test was used for statistical tests. ***: p_value<0.001.

