## Supplementary Table 1-References of lineage-specific marker genes for "Dual-patterned pluripotent stem cells self-organize into a human embryo model with extended anterior-posterior patterning"

**Table S1: References of lineage specific markers.**

| Cell type | Markers used for cell type identification |
| --- | --- |
| MHB[1] | OTX2+[2], EN1+[3], WNT4+[4], PAX2+[5,6], DMBX1+[7] |
| Hindbrain[8] | MAFB+[9], GBX2+[10], HOXA2+[11], HOXD4+[12], PAX6+[13] |
| Hindbrain (r1) | FST+[14], |
| Hindbrain (r2) | FST+[14], CYP26A1+[15], CYP26C1+[16], FGF3+[17] |
| Hindbrain (r3) | EGR2+[18], HOXA2+[11], FGF3+[17] |
| Hindbrain (r4) | HOXB1+[19] |
| Hindbrain (r5) | EGR2+[18], HOXA2+[11], RARB+[20] |
| Hindbrain (r6) | MAFB+[9], RARB+[20] |
| Hindbrain (r7 & r8) | HOXB3+[12], HOXB4+[12], RARB+[20] |
| Roof plate | LMX1A+[21,22], MSX2+[23], WNT1+[24], WNT3A+[25] |
| Neural tube | PAX6+[26], HOXC6+[27], HOXC9+[27] |
| NMPs[28] | TBXT+[29,30], CDX2+[29,31], NKX1-2+[32], SOX2+[33] |
| Neural crest (NC)[34] | SOX10+[35], FOXD3+[36], ETS1+[37] |
| Cranial NC | SPP1+[38], NDST3+[39], NDST4+[39], HOX-)[34] |
| Vagal NC | HOXA3+[40], HOXB3+[40,41], HOXA4+[42], HOXB4+[42] |
| Trunk NC | MSX1+[43], CDX2+[44], NEUROG2+[34] |
| G1 phase | CDKN1A+[45] |
| Early neurons | ONECUT1+[46], ONECUT2+[46] |
| Somitic mesoderm | MEOX1+[47], TCF15+[48] |
| Somites (Differentiation front) | TBX6+[49,50], RIPPLY2+[51], MESP2+[50], MEOX2-[47] |
| Somites (Differentiating somite) | PAX3+[52], FST+[53,54] |
| Somites (Somites) | PAX3+[55] |
| Somites (Myocytes) | NEB+[56,57] |
| Intermediate mesoderm | WT1+[58], OSR1+[59] |
| Renal epithelium | LHX1+[60], PAX2+[61], PAX8+[61] |
| Cardiac mesoderm | HAND2+[62], TNNT2+[63] |
| Head Mesoderm | TWIST1+[34,64], PRRX2+[34,65], FOXC1+[66,67] |
| Gut | SOX17+[68], FOXA2+[69,70] |
| Epiblast | NANOG+[71,72] |

1. Joyner, A. L., Liu, A. & Millet, S. Otx2, Gbx2 and Fgf8 interact to position and maintain a mid-hindbrain organizer. *Current opinion in cell biology* **12**, (2000).
2. Millet, S., Bloch-Gallego, E., Simeone, A. & Alvarado-Mallart, R. M. The caudal limit of Otx2 gene expression as a marker of the midbrain/hindbrain boundary: a study using in situ hybridisation and chick/quail homotopic grafts. *Development* **122**, 3785–3797 (1996).
3. Davis, C. A. & Joyner, A. L. Expression patterns of the homeo box-containing genes En-1 and En-2 and the proto-oncogene int-1 diverge during mouse development. *Genes & development* **2**, (1988).
4. Ungar, A. R., Kelly, G. M. & Moon, R. T. Wnt4 affects morphogenesis when misexpressed in the zebrafish embryo. *Mech Dev* **52**, 153–164 (1995).
5. Urbánek, P., Fetka, I., Meisler, M. H. & Busslinger, M. Cooperation of Pax2 and Pax5 in midbrain and cerebellum development. *Proceedings of the National Academy of Sciences of the United States of America* **94**, (1997).
6. Schwarz, M., Alvarez-Bolado, G., Urbánek, P., Busslinger, M. & Gruss, P. Conserved biological function between Pax-2 and Pax-5 in midbrain and cerebellum development: Evidence from targeted mutations. *Proceedings of the National Academy of Sciences* **94**, 14518–14523 (1997).
7. Dmbx1 is a paired-box containing gene specifically expressed in the caudal most brain structures. *Mechanisms of Development* **114**, 219–223 (2002).
8. Krumlauf, R. & Wilkinson, D. G. Segmentation and patterning of the vertebrate hindbrain. *Development* **148**, (2021).
9. Marín, F. & Charnay, P. Hindbrain patterning: FGFs regulate Krox20 and mafB/kr expression in the otic/preotic region. *Development* **127**, 4925–4935 (2000).
10. Wassarman, K. M. *et al.* Specification of the anterior hindbrain and establishment of a normal mid/hindbrain organizer is dependent on Gbx2 gene function. *Development* **124**, 2923–2934 (1997).
11. Wilkinson, D. G., Bhatt, S., Cook, M., Boncinelli, E. & Krumlauf, R. Segmental expression of Hox-2 homoeobox-containing genes in the developing mouse hindbrain. *Nature* **341**, (1989).
12. Chapter 8 Hox Genes and Segmentation of the Vertebrate Hindbrain. in *Current Topics in Developmental Biology* vol. 88 103–137 (Academic Press, 2009).
13. Kayam, G. *et al.* A novel role for Pax6 in the segmental organization of the hindbrain. *Development (Cambridge, England)* **140**, (2013).
14. Albano, R. M., Arkell, R., Beddington, R. S. P. & Smith, J. C. Expression of inhibin subunits and follistatin during postimplantation mouse development: decidual expression of activin and expression of follistatin in primitive streak, somites and hindbrain. *Development* **120**, 803–813 (1994).
15. MacLean, G. *et al.* Cloning of a novel retinoic-acid metabolizing cytochrome P450, Cyp26B1, and comparative expression analysis with Cyp26A1 during early murine development. *Mech Dev* **107**, 195–201 (2001).
16. Tahayato, A., Dollé, P. & Petkovich, M. Cyp26C1 encodes a novel retinoic acid-metabolizing enzyme expressed in the hindbrain, inner ear, first branchial arch and tooth buds during murine development. *Gene Expr Patterns* **3**, 449–454 (2003).
17. Walshe, J., Maroon, H., McGonnell, I. M., Dickson, C. & Mason, I. Establishment of

hindbrain segmental identity requires signaling by FGF3 and FGF8. *Curr Biol* **12**, 1117–1123 (2002).
18. Wilkinson, D. G., Bhatt, S., Chavrier, P., Bravo, R. & Charnay, P. Segment-specific expression of a zinc-finger gene in the developing nervous system of the mouse. *Nature* **337**, 461–464 (1989).
19. Pöpperl, H. *et al.* Segmental expression of Hoxb-1 is controlled by a highly conserved autoregulatory loop dependent upon exd/pbx. *Cell* **81**, 1031–1042 (1995).
20. Serpente, P. *et al.* Direct crossregulation between retinoic acid receptor β and Hox genes during hindbrain segmentation. *Development* **132**, 503–513 (2005).
21. Chizhikov, V. V. & Millen, K. J. Control of roof plate formation by Lmx1a in the developing spinal cord. *Development* **131**, 2693–2705 (2004).
22. Millonig, J. H., Millen, K. J. & Hatten, M. E. The mouse Dreher gene Lmx1a controls formation of the roof plate in the vertebrate CNS. *Nature* **403**, (2000).
23. Duval, N. *et al.* Msx1 and Msx2 act as essential activators of Atoh1 expression in the murine spinal cord. *Development* **141**, 1726–1736 (2014).
24. Megason, S. G. & McMahon, A. P. A mitogen gradient of dorsal midline Wnts organizes growth in the CNS. *Development* **129**, 2087–2098 (2002).
25. Roelink, H. & Nusse, R. Expression of two members of the Wnt family during mouse development--restricted temporal and spatial patterns in the developing neural tube. *Genes & development* **5**, (1991).
26. Ericson, J. *et al.* Pax6 controls progenitor cell identity and neuronal fate in response to graded Shh signaling. *Cell* **90**, (1997).
27. Coughlan, E. *et al.* A Hox Code Defines Spinocerebellar Neuron Subtype Regionalization. *Cell Rep* **29**, 2408–2421.e4 (2019).
28. Henrique, D., Abranches, E., Verrier, L. & Storey, K. G. Neuromesodermal progenitors and the making of the spinal cord. *Development* **142**, 2864–2875 (2015).
29. Amin, S. *et al.* Cdx and T Brachyury Co-activate Growth Signaling in the Embryonic Axial Progenitor Niche. *Cell Rep* **17**, 3165–3177 (2016).
30. Gouti, M. *et al.* In vitro generation of neuromesodermal progenitors reveals distinct roles for wnt signalling in the specification of spinal cord and paraxial mesoderm identity. *PLoS biology* **12**, (2014).
31. Savory, J. G. A. *et al.* Cdx2 regulation of posterior development through non-Hox targets. *Development* **136**, 4099–4110 (2009).
32. Schubert, F. R., Fainsod, A., Gruenbaum, Y. & Gruss, P. Expression of the novel murine homeobox gene Sax-1 in the developing nervous system. *Mech Dev* **51**, 99–114 (1995).
33. Graham, V., Khudyakov, J., Ellis, P. & Pevny, L. SOX2 functions to maintain neural progenitor identity. *Neuron* **39**, 749–765 (2003).
34. Soldatov, R. *et al.* Spatiotemporal structure of cell fate decisions in murine neural crest. *Science (New York, N.Y.)* **364**, (2019).
35. Southard-Smith, E. M., Kos, L. & Pavan, W. J. Sox10 mutation disrupts neural crest development in Dom Hirschsprung mouse model. *Nature genetics* **18**, (1998).
36. Dottori, M., Gross, M. K., Labosky, P. & Goulding, M. The winged-helix transcription factor Foxd3 suppresses interneuron differentiation and promotes neural crest cell fate. *Development (Cambridge, England)* **128**, (2001).

37. Théveneau, E., Duband, J. L. & Altabef, M. Ets-1 confers cranial features on neural crest delamination. *PloS one* **2**, (2007).
38. Thayer, J. M. & Schoenwolf, G. C. Early expression of Osteopontin in the chick is restricted to rhombomeres 5 and 6 and to a subpopulation of neural crest cells that arise from these segments. *Anat Rec* **250**, 199–209 (1998).
39. Heparan sulfate expression in the neural crest is essential for mouse cardiogenesis. *Matrix Biology* **35**, 253–265 (2014).
40. Gogolou, A. *et al.* Early anteroposterior regionalisation of human neural crest is shaped by a pro-mesodermal factor. *Elife* **11**, (2022).
41. Chan, K. K. *et al.* Hoxb3 vagal neural crest-specific enhancer element for controlling enteric nervous system development. *Dev Dyn* **233**, 473–483 (2005).
42. Pitera, J. E., Smith, V. V., Thorogood, P. & Milla, P. J. Coordinated expression of 3' hox genes during murine embryonal gut development: an enteric Hox code. *Gastroenterology* **117**, 1339–1351 (1999).
43. Frith, T., Jr *et al.* Human axial progenitors generate trunk neural crest cells in vitro. *Elife* **7**, (2018).
44. Hackland, J. O. S. *et al.* FGF Modulates the Axial Identity of Trunk hPSC-Derived Neural Crest but Not the Cranial-Trunk Decision. *Stem Cell Reports* **12**, 920–933 (2019).
45. Sherr, C. J. & Roberts, J. M. CDK inhibitors: positive and negative regulators of G1-phase progression. *Genes Dev* **13**, 1501–1512 (1999).
46. van der Raadt, J., van Gestel, S. H. C., Nadif Kasri, N. & Albers, C. A. ONECUT transcription factors induce neuronal characteristics and remodel chromatin accessibility. *Nucleic Acids Res* **47**, 5587–5602 (2019).
47. Reijntjes, S., Stricker, S. & Mankoo, B. S. A comparative analysis of Meox1 and Meox2 in the developing somites and limbs of the chick embryo. *Int J Dev Biol* **51**, 753–759 (2007).
48. Burgess, R., Rawls, A., Brown, D., Bradley, A. & Olson, E. N. Requirement of the paraxis gene for somite formation and musculoskeletal patterning. *Nature* **384**, 570–573 (1996).
49. White, P. H., Farkas, D. R., McFadden, E. E. & Chapman, D. L. Defective somite patterning in mouse embryos with reduced levels of Tbx6. *Development* **130**, 1681–1690 (2003).
50. Oginuma, M., Niwa, Y., Chapman, D. L. & Saga, Y. Mesp2 and Tbx6 cooperatively create periodic patterns coupled with the clock machinery during mouse somitogenesis. *Development (Cambridge, England)* **135**, (2008).
51. Biris, K. K., Dunty, W. C., Jr & Yamaguchi, T. P. Mouse Ripply2 is downstream of Wnt3a and is dynamically expressed during somitogenesis. *Dev Dyn* **236**, 3167–3172 (2007).
52. Wiggan, O. 'neil, Fadel, M. P. & Hamel, P. A. Pax3 induces cell aggregation and regulates phenotypic mesenchymal-epithelial interconversion. *J Cell Sci* **115**, 517–529 (2002).
53. Connolly, D. J., Patel, K., Seleiro, E. A., Wilkinson, D. G. & Cooke, J. Cloning, sequencing, and expressional analysis of the chick homologue of follistatin. *Dev Genet* **17**, 65–77 (1995).
54. Monica, S. D. & Harland, R. M. Follistatin interacts with Noggin in the development of the axial skeleton. *Mechanisms of development* **131**, (2014).
55. Goulding, M., Lumsden, A. & Paquette, A. J. Regulation of Pax-3 expression in the dermomyotome and its role in muscle development. *Development* **120**, 957–971 (1994).
56. Labeit, S. & Kolmerer, B. The complete primary structure of human nebulin and its

correlation to muscle structure. *J Mol Biol* **248**, 308–315 (1995).
57. Kruger, M., Wright, J. & Wang, K. Nebulin as a length regulator of thin filaments of vertebrate skeletal muscles: correlation of thin filament length, nebulin size, and epitope profile. *J Cell Biol* **115**, 97–107 (1991).
58. Armstrong, J. F., Pritchard-Jones, K., Bickmore, W. A., Hastie, N. D. & Bard, J. B. The expression of the Wilms' tumour gene, WT1, in the developing mammalian embryo. *Mechanisms of development* **40**, (1993).
59. Mugford, J. W., Sipilä, P., McMahon, J. A. & McMahon, A. P. Osr1 expression demarcates a multi-potent population of intermediate mesoderm that undergoes progressive restriction to an Osr1-dependent nephron progenitor compartment within the mammalian kidney. *Developmental biology* **324**, (2008).
60. Cirio, M. C. *et al.* Lhx1 is required for specification of the renal progenitor cell field. *PloS one* **6**, (2011).
61. Bouchard, M., Souabni, A., Mandler, M., Neubüser, A. & Busslinger, M. Nephric lineage specification by Pax2 and Pax8. *Genes & development* **16**, (2002).
62. Tsuchihashi, T. *et al.* Hand2 function in second heart field progenitors is essential for cardiogenesis. *Developmental biology* **351**, (2011).
63. England, J., Pang, K. L., Parnall, M., Haig, M. I. & Loughna, S. Cardiac troponin T is necessary for normal development in the embryonic chick heart. *Journal of anatomy* **229**, (2016).
64. Bildsoe, H. *et al.* Transcriptional targets of TWIST1 in the cranial mesoderm regulate cell-matrix interactions and mesenchyme maintenance. *Developmental biology* **418**, (2016).
65. Higuchi, M. *et al.* Temporospatial gene expression of Prx1 and Prx2 is involved in morphogenesis of cranial placode-derived tissues through epithelio-mesenchymal interaction during rat embryogenesis. *Cell and tissue research* **353**, (2013).
66. Mya, N. *et al.* Transcription factor Foxc1 is involved in anterior part of cranial base formation. *Congenital Anomalies* **58**, 158–166 (2018).
67. Mittnenzweig, M. *et al.* A single-embryo, single-cell time-resolved model for mouse gastrulation. *Cell* **184**, (2021).
68. Kanai-Azuma, M. *et al.* Depletion of definitive gut endoderm in Sox17-null mutant mice. *Development* **129**, 2367–2379 (2002).
69. Sasaki, H. & Hogan, B. L. Differential expression of multiple fork head related genes during gastrulation and axial pattern formation in the mouse embryo. *Development* **118**, 47–59 (1993).
70. Monaghan, A. P., Kaestner, K. H., Grau, E. & Schütz, G. Postimplantation expression patterns indicate a role for the mouse forkhead/HNF-3 alpha, beta and gamma genes in determination of the definitive endoderm, chordamesoderm and neuroectoderm. *Development* **119**, 567–578 (1993).
71. Loh, Y.-H. *et al.* The Oct4 and Nanog transcription network regulates pluripotency in mouse embryonic stem cells. *Nature Genetics* **38**, 431–440 (2006).
72. Silva, J. *et al.* Nanog is the gateway to the pluripotent ground state. *Cell* **138**, (2009).
